# Protein arginine methyltransferase 5 promotes metastasis via enhancing EGFR transcription and modulating AKT1 activation by methylation

**DOI:** 10.1101/2020.08.12.246660

**Authors:** Lei Huang, Xiao-Ou Zhang, Odette Verdejo-Torres, Kim Wigglesworth, Xiaomei Sun, Benjamin Sallis, Daniel Moon, Tingting Huang, Esteban Rozen, Gang Wang, Lei Zhang, Jason Shohet, Mary M. Lee, Qiong Wu

## Abstract

Protein arginine methyltransferase 5 (PRMT5) regulates a wide range of physiological processes, including cancer cell proliferation and metastasis, by generating symmetric di-methyl-arginine marks on histones and non-histone proteins. Here, we report that PRMT5 directly regulates epidermal growth factor receptor (EGFR) transcription to control EGF stimulated EGFR signaling. Furthermore, PRMT5 modulates protein kinase B (AKT) activation by methylation of AKT1 Arg 15, which is required for its subsequent phosphorylation at AKT1 Thr 308 and Ser 473. The PRMT5/EGFR/AKT axis converges to regulate transcription factors ZEB1, SNAIL, and TWIST1 to promote the epithelial-mesenchymal transition (EMT), in the manner that EGFR and AKT1 compensate each other to support tumor cell invasion and metastasis. Inhibiting PRMT5 methyltransferase activity with a small molecule inhibitor attenuated primary tumor growth and prevented hepatic metastasis in aggressive *in vivo* tumor models. Collectively, our results support the use of PRMT5 based therapies for metastatic cancer.

## Introduction

Epithelial-mesenchymal transition (EMT) plays an essential role in the formation of tissues and organs during embryogenesis and is vital in the repair of damaged tissue in adults^1–3^. However, dysregulated EMT also confers tumor cell stemness, invasiveness, and drug resistance^4–7^. A group of transcription factors (TFs), including SNAIL, ZEB, and TWIST, orchestrate the EMT program ^8, 9^. The EMT-TFs are induced by extracellular stimuli, including TGF-β, growth factors via receptor tyrosine kinases (RTK) such as EGFR and FGFR, paracrine signaling through Notch ligand and Wnt, and the hypoxic tumor microenvironment and tumor-associated stroma^10–12^. On the molecular level, the EMT process is modulated by multiple signaling pathways, some of which are mediated by phosphoinositide 3-kinase (PI3K)/protein kinase B (AKT)/mammalian target of rapamycin (mTOR)^13–15^.

Contrary to tumor initiation, which is typically driven by somatic mutations or translocations, tumor cell invasion and metastasis are mainly driven by non-genetic processes, such as EMT and epigenetic mechanisms. Epigenetic regulators such as the polycomb repressive complexes (PRCs), histone acetyltransferases (HATs) and deacetylases (HDACs), histone methyltransferases (HMTs) and demethylases (HDMs), DNA methyltransferases (DNMTs), nucleosome remodelers and in particular, protein arginine methyltransferase 5 (PRMT5), have been shown to participate by regulating EMT-TFs and EMT markers^16, 17^. For example, the PRMT5-MEP50 complex methylates H3R2 to induce EMT-TFs activator genes; and also methylates H4R3 to repress metastasis suppressor genes E-cadherin and GAS1 in lung and breast cancer^18^.

Neuroblastoma is a neural crest-derived embryonal malignancy in infants^19, 20^. Early embryonic neural crest precursors undergo a spatially and temporally programmed EMT and differentiation process as they migrate out of the neural tube away from the primitive neural tubes and contribute to the formation of over seventy different tissue types. A subset of ventral neural crest precursors is destined to form the paraspinal sympathetic ganglia and adrenal glands from which neuroblastomas in infants and young children arise^20–24^. Inhibition of programmed differentiation is thought to drive tumor formation^20^. In epithelial cancers, it is well accepted that EMT supports cellular invasiveness, a major contributor to metastatic potential^7, 25^. We hypothesized that the persistence of the EMT program beyond the early physiological spatial patterns and temporal windows in neural crest precursors contribute to the pathogenesis of highly metastatic and aggressive neuroblastomas^20^. During neural crest development, AKT signaling regulates EMT by multiple mechanisms but also has been implicated in cancer-specific EMT, invasion, and metastasis^21, 26–28^. As such, combinational targeting of the PI3K/AKT/mTOR signaling networks has been extensively explored as a therapeutic approach to limit or block tumor metastasis^29–32^.

Protein arginine methyltransferase 5 (PRMT5), a member of type II arginine methyltransferases, is the primary methyltransferase generating symmetric dimethylarginine (SDMA)^33–35^. Arginine methylation plays a critical role in transcriptional regulation through histone arginine methylation, but also via modification of glycine-arginine rich motifs RGG/RG) involved in protein/DNA and protein/protein interactions^35, 36^. We have previously shown that PRMT5 increases the expression of the adipogenic master transcriptional factor PPARγ2 during pre-adipocyte differentiation^37, 38^, and elevates AKT2 activation in hepatocytes in response to dietary fatty acid stimulation^39^. Our initial findings suggested the clinical relevance of PRMT5 in neuroblastoma. We report now that PRMT5 inhibition by a small molecule inhibitor or shRNA knockdown markedly decreases cell proliferation in cultured neuroblastoma cells and attenuates tumor growth in xenograft models. Most critically, a PRMT5 inhibitor blocks tumor cell metastasis to the liver, a common metastatic site for neuroblastomas. We find that PRMT5 transcriptionally regulates EGFR signaling and post-translationally modulates AKT1 activation via methylation. We demonstrate that PRMT5 mediated arginine methylation is required for AKT1 activation. Furthermore, we show that PRMT5 directly and independently regulates EGFR and AKT signaling, which converge to regulate the expression of EMT-TFs ZEB, TWIST1, and SNAIL through which EMT networks are directed to promote PRMT5 mediated tumor metastasis. Thus, using both *in vitro* and *in vivo* approaches, we define a central role of PRMT5 in EMT and the underlying mechanisms by which it regulates the process of metastasis. PRMT5 inhibition may represent a novel therapeutic tool to prevent metastatic cancers.

## Results

### PRMT5 is dysregulated in advanced stage neuroblastoma

To explore the clinical significance of PRMT5, we analyzed its expression in the Tumor Alterations Relevant for Genomics-driven Therapy (TARGET) database containing RNA-seq results from 161 patients with stage 3 and stage 4 neuroblastoma in which tumors spread to regional lymph nodes or distal lymph nodes, respectively. PRMT5 transcript levels were higher in stage 4 patients than stage 3 patients (Fig. 1a, left). Importantly, high levels of PRMT5 correlated with worse survival (Fig. 1a, right). We also performed this analysis in three additional independent neuroblastoma annotated patient cohorts available in the R2 database (R2: Genomics Analysis and Visualization Platform(http://r2.amc.nl)). We found that PRMT5 transcripts were more abundant in stage 4 than stage 3 patients and negatively correlated with patient survival (Fig. 1b-d).

**Figure 1.**
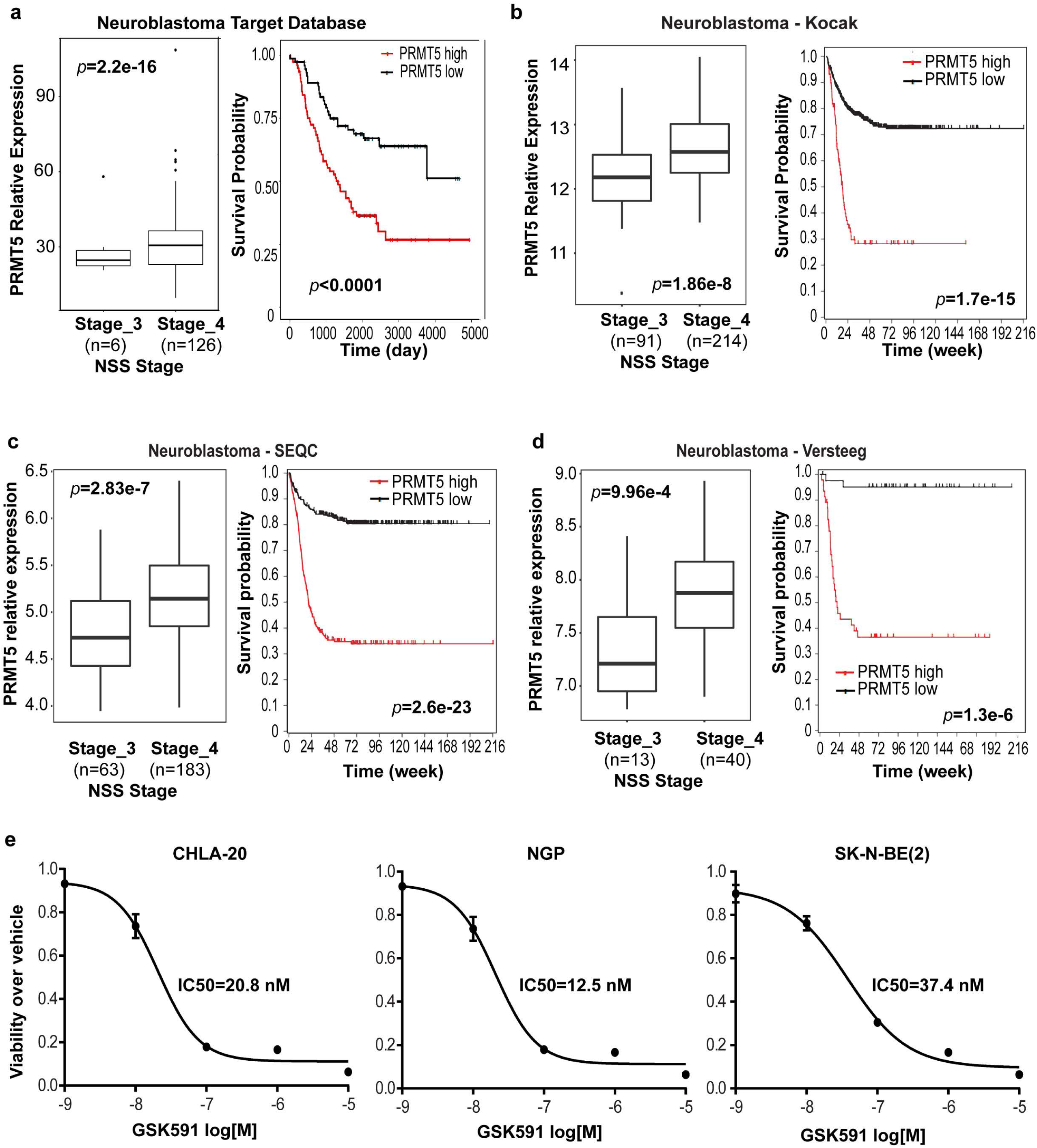
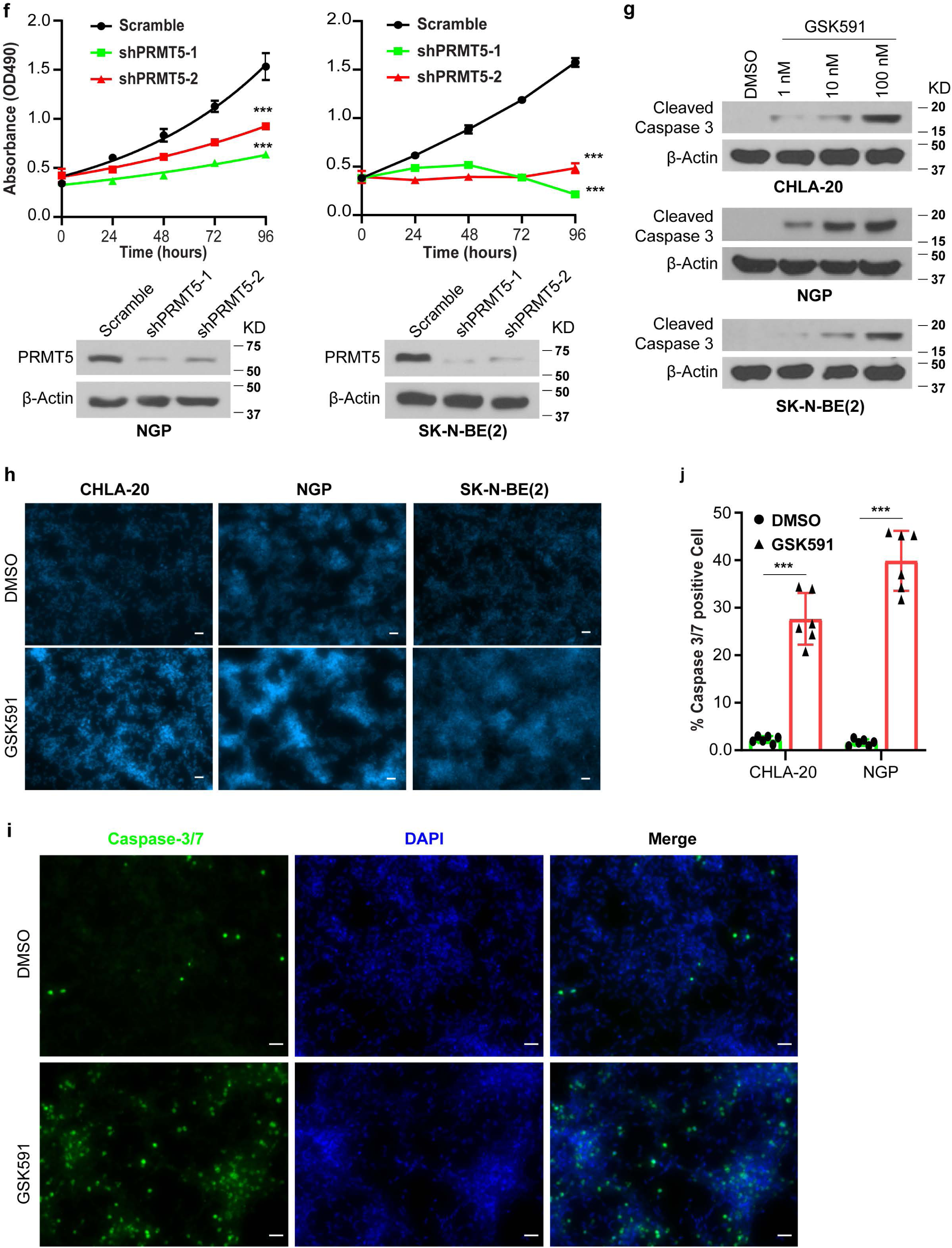
Overexpression of PRMT5 is associated with high-risk neuroblastoma and poor patient survival and is required for neuroblastoma cell proliferation *in vitro*. **a**, The relative expression of PRMT5 (right) and correlation of PRM5 levels with patient survival probability (left) by analysis of RNA-seq data from 161 neuroblastoma patients in the TARGET database. PRMT5 expression levels (left) and patient survival probability (right) in stage 3 and stage 4 neuroblastoma patients in three other databases, as shown in **b**, **c** and **d**. **e**. Wilcoxon rank-sum test *p* values for boxplots (left) and log-rank test *p* values for survival curves (right) are shown in **a**-**d**. The efficacy of PRMT5 small molecule inhibitor GSK3203591 (GSK591) determined by MTS assay in CHLA20, NGP, and SK-N-BE (2) cells. **f**, Cell viability measured by MTA assay in a scramble or shRNA targeting PRMT5 in NGP (left) and SK-N-BE (2) cells (right) (*n* = 3). **g**, Cleaved caspase 3 levels in neuroblastoma cell lines treated with increasing doses of GSK591. **h**, Hoechst 33342 staining in cells treated with DMSO or 100 nM GSK591. Scale bars, 100 μm. **i**, Apoptosis measured by caspase-3/7 staining in CHLA20 cells treated with DMSO or GSK591. **j**, Quantification of caspase-3/7 positive cells. ***, *p*<0.0001. Statistics were analyzed by the two-tailed *t*-test in GraphPad Prism (**e**, **f**, **j**). **e**-**i**, three independent experiments.

### PRMT5 inhibition reduces neuroblastoma cell viability

Given its involvement in many cancers, PRMT5 has become a promising target for therapeutic intervention^19, 35, 40^. Two small molecule inhibitors targeting PRMT5 methyltransferase activity, GSK3326595 (clinical trial identifier NCT02783300) and JNJ-64619178 (clinical trial identifier NCT03573310), have been or are currently being tested in early-phase clinical trials for the treatment of advanced solid tumors, acute myeloid leukemia, and non-Hodgkin’s lymphoma. For *in vivo* studies, we used GSK3326595 (GSK595, hereafter) due to its confirmed efficacy in a breast cancer mouse model with favorable *in vivo* PD/PK; for *in vitro* studies, we used its sister compound GSK3302591 (GSK591, hereafter) that shows increased cell permeability in tissue culture. We tested these PRMT5 inhibitors in three neuroblastoma cell lines representative of the clinical diversity of metastatic neuroblastoma, 1) having high c-MYC level without N-MYC amplification and derived from a relapsed tumor (CHLA20), 2) having low c-MYC with N-MYC amplification and derived from a pretreatment tumor (NGP) and 3) from a relapsed tumor (SK-N-BE(2)). As shown in Fig. 1e, PRMT5 inhibition by GSK591 in the low nanomolar range significantly decreased neuroblastoma cell viability. PRMT5 generates the majority of cellular symmetric dimethylarginine (SDMA); thus, we examine the global SDMA as a readout of PRMT5 enzymatic activity. We verified that GSK591 treatment significantly decreased SDMA in a dose-dependent manner, indicating the on-target effects of this compound (Supplemental Fig. 1a). We generated doxycycline-inducible shRNA mediated PRMT5 knockdown cell lines to confirm these results (Supplemental Fig. 1b). Again, induced depletion of PRMT5 decreased global SDMA (Supplemental Fig. 1c) and neuroblastoma cell viability (Fig. 1f). In agreement with the reduced cell viability, we found that GSK 591 treatment triggered apoptosis in a dose-dependent manner, as evidenced by the increasing levels of cleaved caspase 3 (Fig. 1g). Also, more condensed, pycnotic nuclei were seen in GSK591 treated cells than in control cells, as visualized by Hoechst 33342 (Fig. 1h). When stained with a green fluorescent dye recognizing cleaved caspase-3/7, GSK591 treated cells had a higher percentage of caspase-3/7 positive cells (Fig. 1i, j, Supplemental Fig. 1d).

### PRMT5 inhibition attenuates primary tumor growth and blocks hepatic metastasis

We then tested the effects of GSK595 in an orthotopic mouse xenograft model. To monitor the tumor growth by live animal imaging, we transduced CHLA20 or NGP cells with iRFP 720-Luc reporter lentiviral vectors that allowed *in vivo* imaging by bioluminescence and differentiated human tumor cells from mouse cells by iRFP 720. Successfully transduced cells were enriched by FACS sorting on the iRFP 720 signal. We implanted neuroblastoma cells into the renal capsule of NOD/SCID mice. This is a well-characterized model that faithfully recapitulates the aggressive, highly vascular growth neuroblastoma from the adrenal parasympathetic precursors, as well as local invasiveness and metastatic spread of human neuroblastoma^41^. Animals were treated for two weeks with 100 mg/kg GSK595 or with vehicle by oral gavage twice daily (Fig. 2a). Serum blood chemistries showed no significant toxicity (Table 2), and there was no significant weight loss after two weeks of GSK595 treatment (Supplemental Fig. 2a). The effective delivery of GSK595 to xenograft tumors was confirmed by examining SDMA levels in tumor tissues from control and GSK595 treated mice (Supplemental Fig. 2b, c). The tumor mass from the GSK595 treated group was significantly reduced in xenografts of CHLA20 (Fig. 2b) and NGP (Fig. 2e) compared to control mice. Light microscopy showed that the size of the *ex vivo* tumors from the vehicle-treated mice was much larger than that from GSK595 treated mice in both CHLA20 and NGP xenografts, respectively (Fig. 2c, f). Bioluminescent imaging demonstrated that there were fewer tumor cells in the GSK595 treated tumor tissues compared to vehicle-treated mice (Fig. 2d, g). Importantly, we found fewer tumor cells in the livers from mice treated with GSK595 than in control mice by bioluminescent imaging (Fig. 2h). This observation was confirmed by FACS analysis of iRFP 720 positive human neuroblastoma cells that were implanted (Fig. 2i, j). Collectively, GSK595 treatment suppressed primary tumor growth and blocked tumor cell metastasis to the liver.

**Figure 2.**
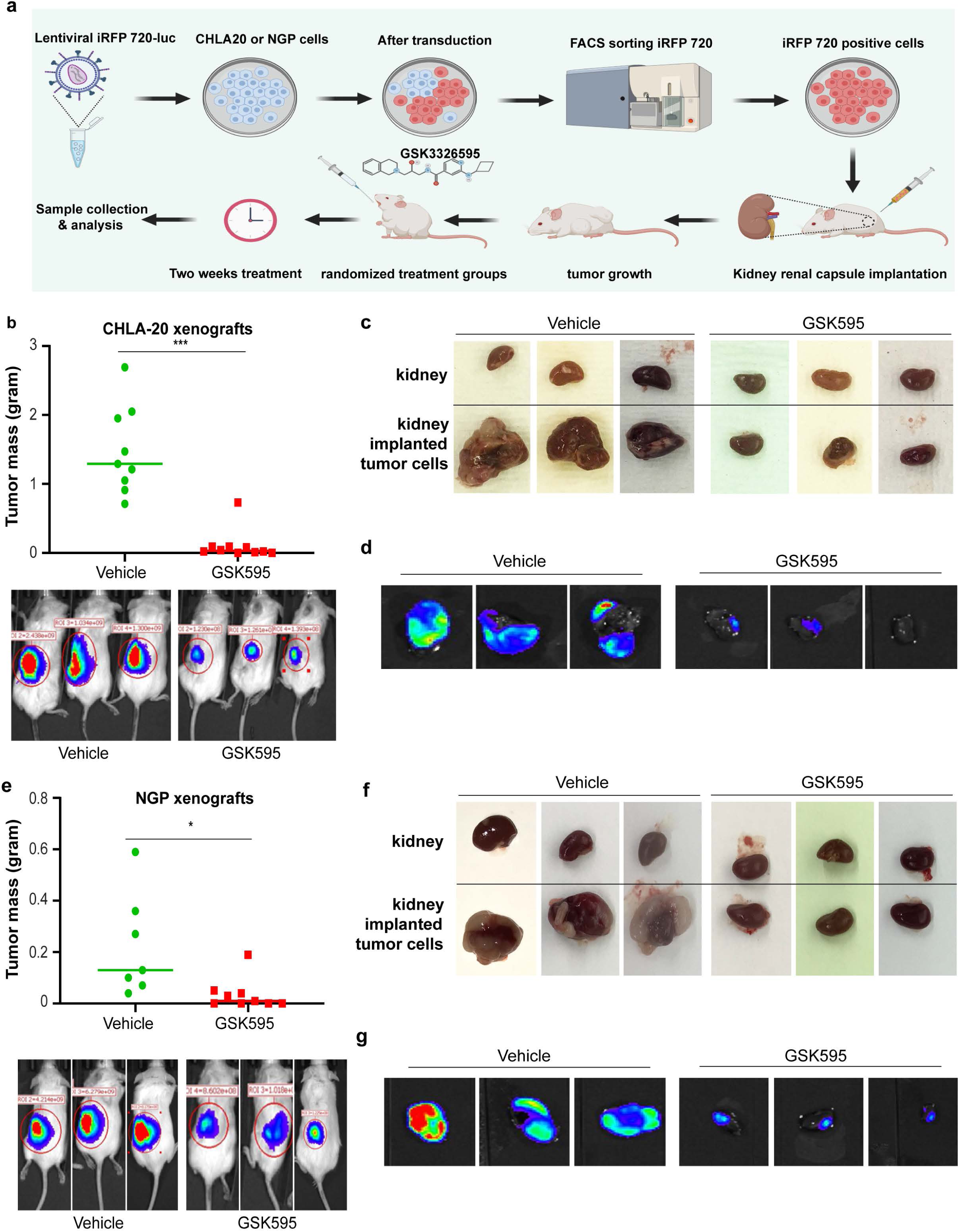
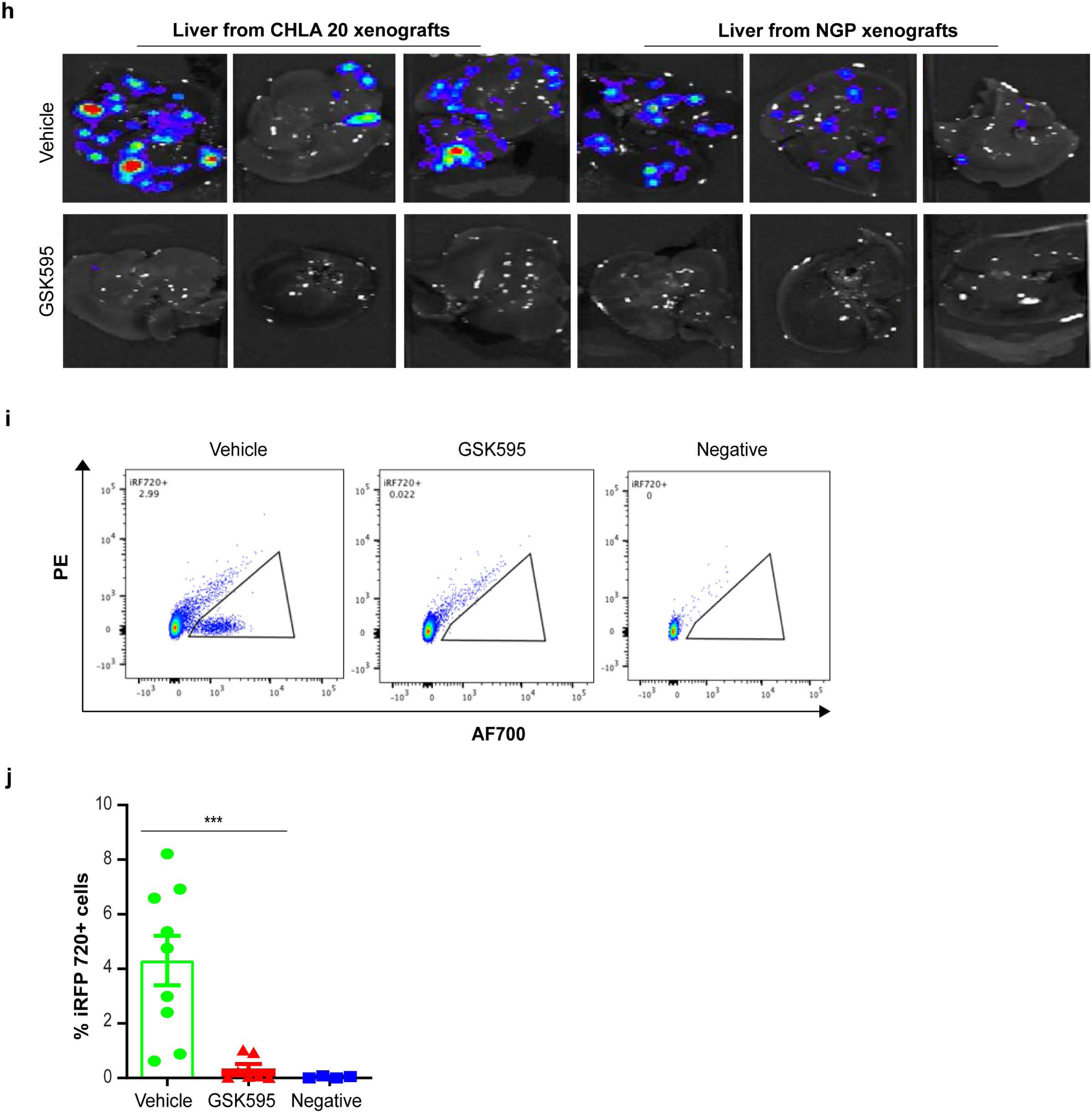
Inhibiting PRMT5 methyltransferase activity attenuated primary tumor growth and metastasis. **a**, Schematic of PRMT5 inhibitor GSK3326595 (GSK595) *in vivo* study in the kidney renal capsule implantation xenograft model. Neuroblastoma cells were transduced with lentiviral iRFP 720-luc and iRFP 720 positive cells were enriched by fluorescence-activated cell sorting (FACS). One million iRFP 720-Luc cells were injected into the renal capsule of the right kidney, and tumor growth was monitored by live-animal bioluminescent imaging. Once tumors reached to 10^7^∼10^8^ average radiance by bioluminescence, animals were randomized to vehicle or GSK595 groups. Mice were treated twice daily with 100 mg/kg GSK595 by oral gavage for two weeks. **b**, Tumor mass of CHLA20 iRFP 720-Luc xenograft tumors from mice treated with vehicle or GSK595 (n=9-10) (top) and representative *in vivo* bioluminescent image and quantification of tumor (bottom). Tumor mass was calculated by subtracting the weight of the normal kidney from the weight of the kidney that was implanted in tumor cells. Representative *ex vivo* images of tumor by light microscope (**c**) or bioluminescence (**d**). **e**, Tumor mass of NGP iRFP 720-Luc xenograft tumors from mice treated with vehicle or GSK595 (n=7-9) (top) and representative *in vivo* bioluminescent images and quantification of tumor (bottom). Representative *ex vivo* images of tumor by light microscope (**f**) or bioluminescence (**g**). **h**, Representative *ex vivo* bioluminescent images of liver from mice bearing CHLA 20 or NGP xenograft tumors treated with vehicle or GSK595. **i**, Representative charts of FACS analysis of iRFP720 positive human neuroblastoma cells in hepatocytes isolated from the whole liver from CHHLA20 xenografted mice. **j**, Quantification of the percentage of iRFP 720 positive tumor cells determined by flow cytometry in hepatocytes isolated from the whole liver from CHHLA20 xenografted mice. ***, *p*<0.0001. Statistics were analyzed by two-tailed t test in GraphPad Prism.

**Table 1.**
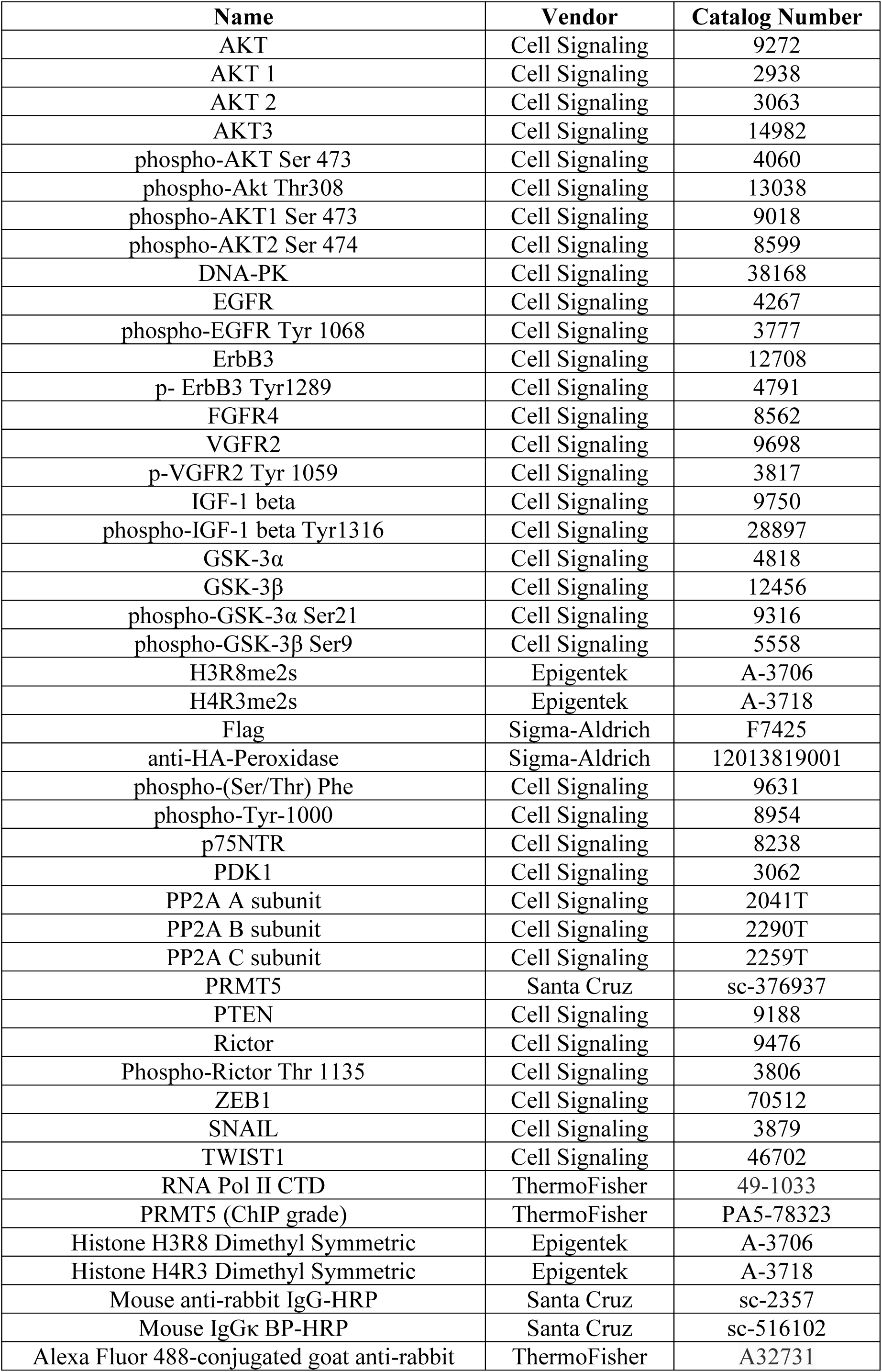
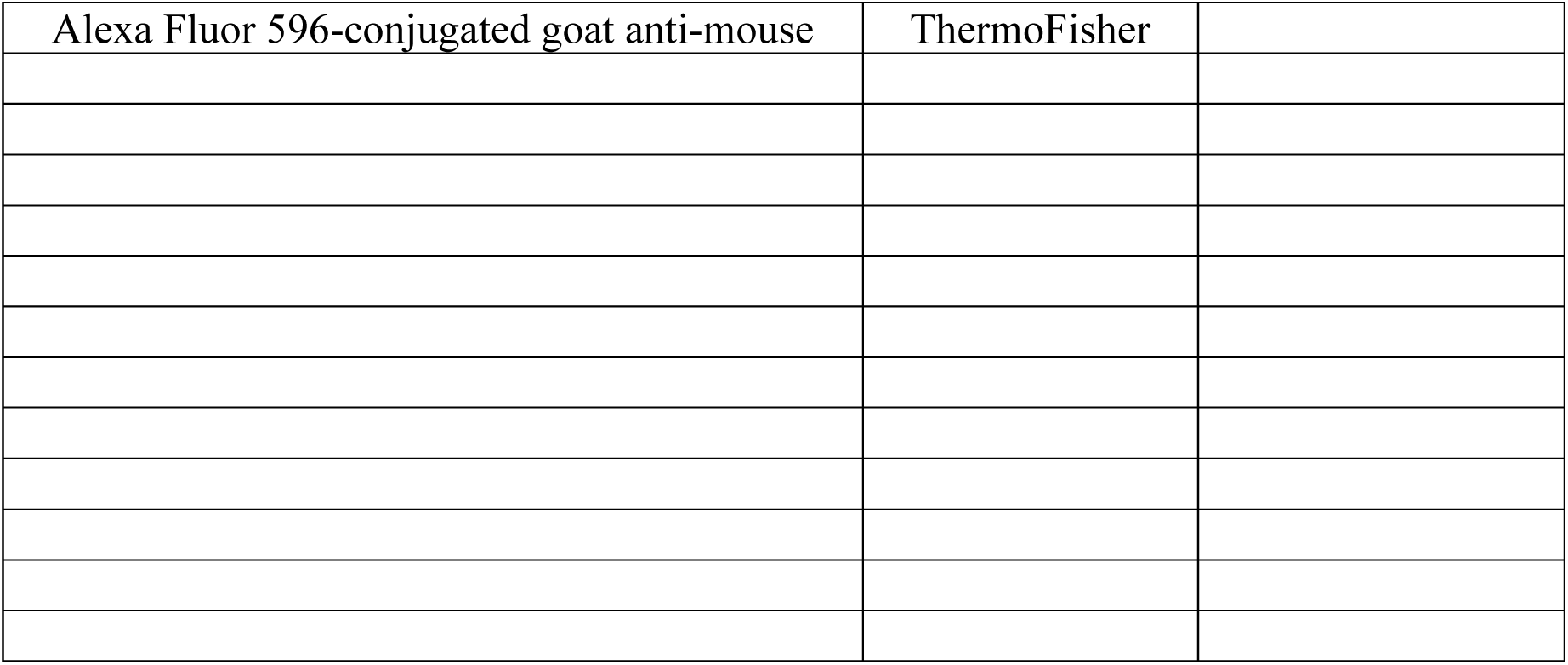
Antibodies used in this study.

**Table 2.**
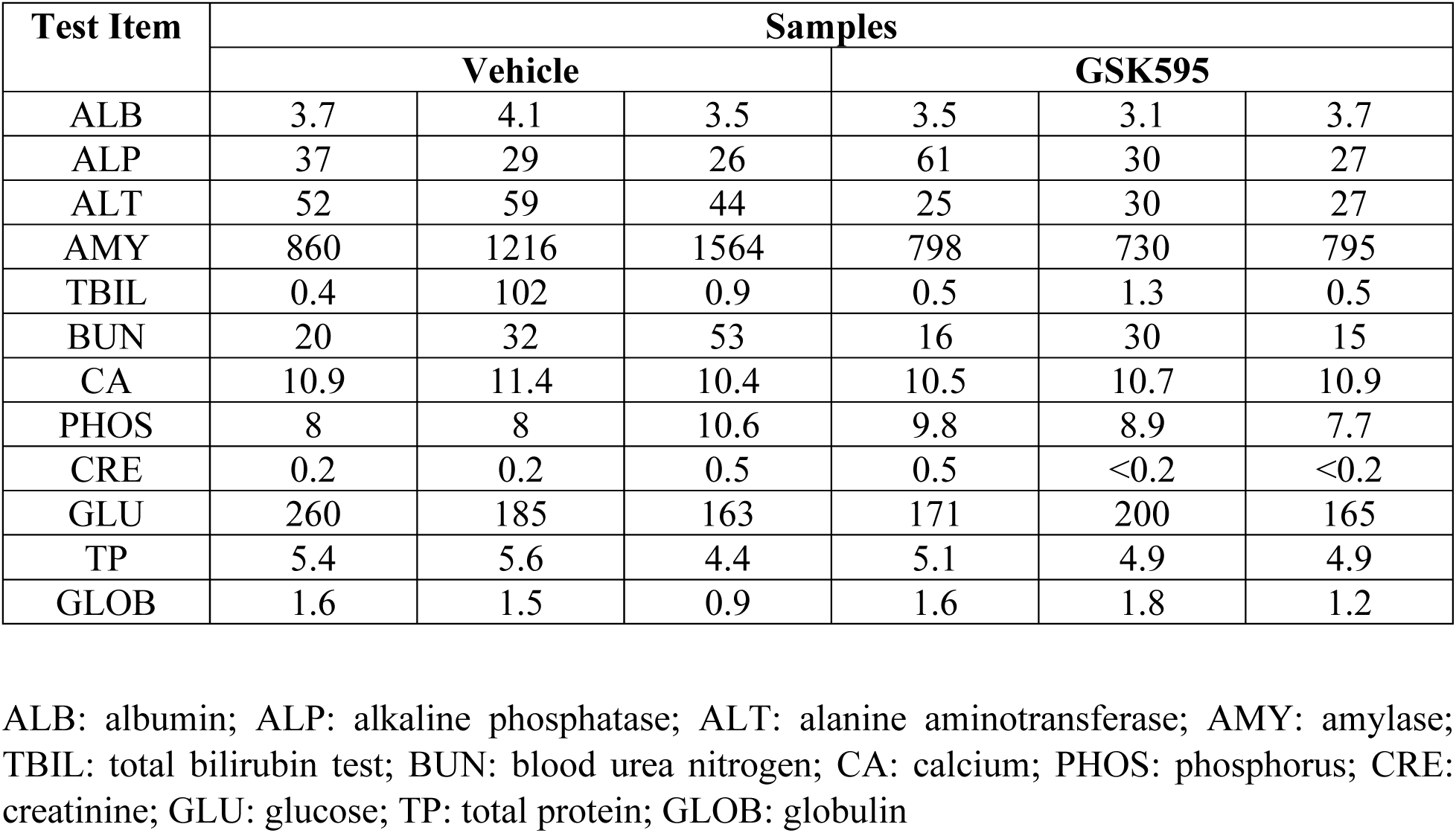
Mouse Blood Chemistry Test.

| Test Item | Samples |  |  |  |  |  |
| --- | --- | --- | --- | --- | --- | --- |
|  | Vehicle |  |  | GSK595 |  |  |
| ALB | 3.7 | 4.1 | 3.5 | 3.5 | 3.1 | 3.7 |
| ALP | 37 | 29 | 26 | 61 | 30 | 27 |
| ALT | 52 | 59 | 44 | 25 | 30 | 27 |
| AMY | 860 | 1216 | 1564 | 798 | 730 | 795 |
| TBIL | 0.4 | 102 | 0.9 | 0.5 | 1.3 | 0.5 |
| BUN | 20 | 32 | 53 | 16 | 30 | 15 |
| CA | 10.9 | 11.4 | 10.4 | 10.5 | 10.7 | 10.9 |
| PHOS | 8 | 8 | 10.6 | 9.8 | 8.9 | 7.7 |
| CRE | 0.2 | 0.2 | 0.5 | 0.5 | <0.2 | <0.2 |
| GLU | 260 | 185 | 163 | 171 | 200 | 165 |
| TP | 5.4 | 5.6 | 4.4 | 5.1 | 4.9 | 4.9 |
| GLOB | 1.6 | 1.5 | 0.9 | 1.6 | 1.8 | 1.2 |

ALB: albumin; ALP: alkaline phosphatase; ALT: alanine aminotransferase; AMY: amylase; TBIL: total bilirubin test; BUN: blood urea nitrogen; CA: calcium; PHOS: phosphorus; CRE: creatinine; GLU: glucose; TP: total protein; GLOB: globulin

**Table 3.**
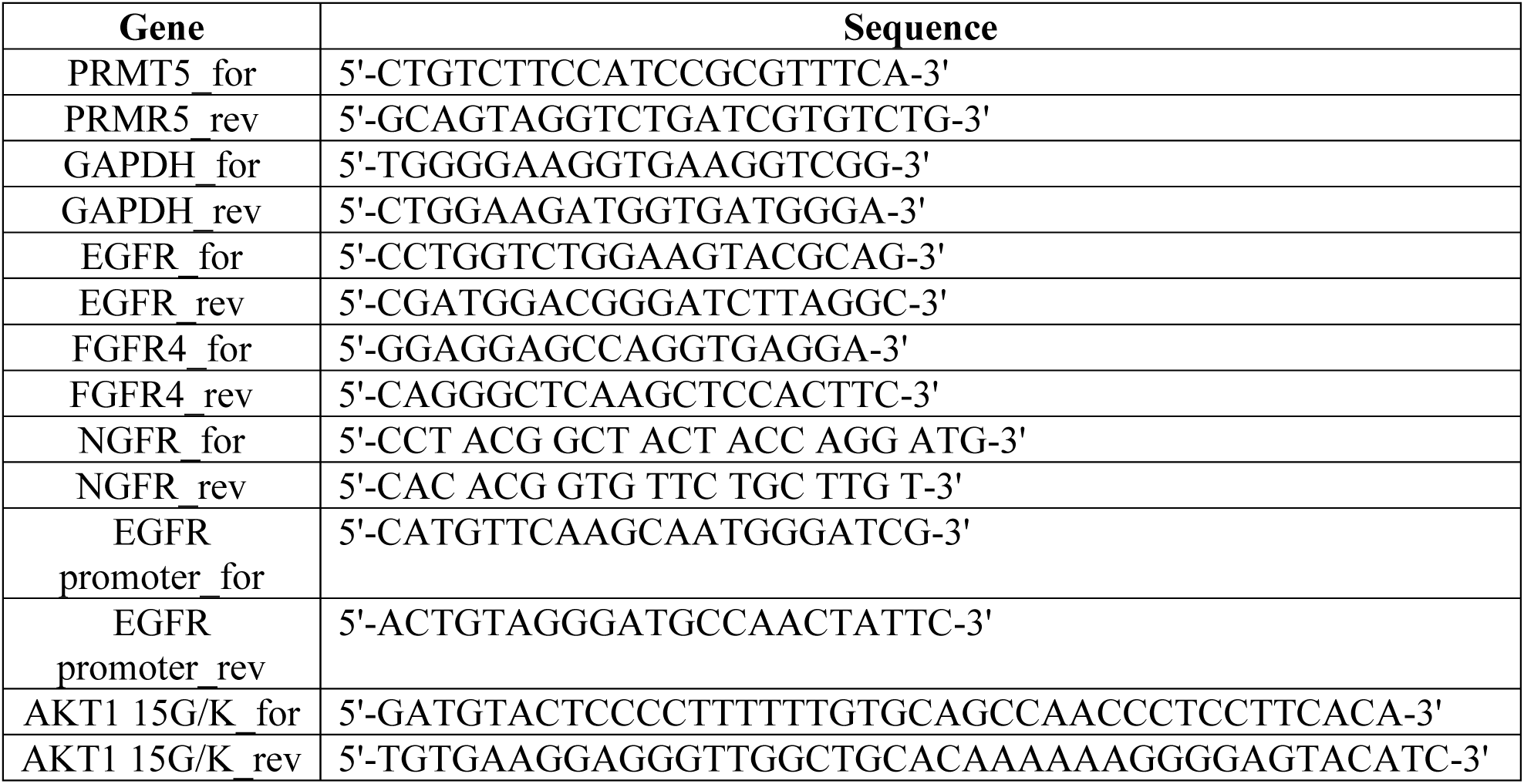
Primers used in this study.

### Global transcriptome analysis in neuroblastoma cells after PRMT5 inhibition

We investigated the mechanisms by which PRMT5 regulates neuroblastoma gene expression by RNA-seq. Changes in global gene expression were identified in CHLA20 and NGP cells treated with DMSO or GSK591 by unsupervised hierarchical clustering (Fig. 3a). There was a small subset of overlap in the differentially expressed genes (DEGs) in these two cell lines, which may indicate the different gene networks mediated by c-MYC (CHLA20) compared to N-MYC (NGP), respectively (Fig. 3b). The top pathways in KEGG (Kyoto Encyclopedia of Gene and Genomes) pathway analysis (Fig. 3c, d) showed that the downregulated DEGs were mostly enriched in PI3K-AKT and p53 signaling pathways in both cell lines. In gene set enrichment analysis, the GSK591 treated CHLA20 cells had a decrease in the AKT up- and down-regulated genes (Fig. 3e), whereas in NGP cells, GSK591 treatment downregulated AKT/mTOR specific genes (Fig. 3f).

**Figure 3.**
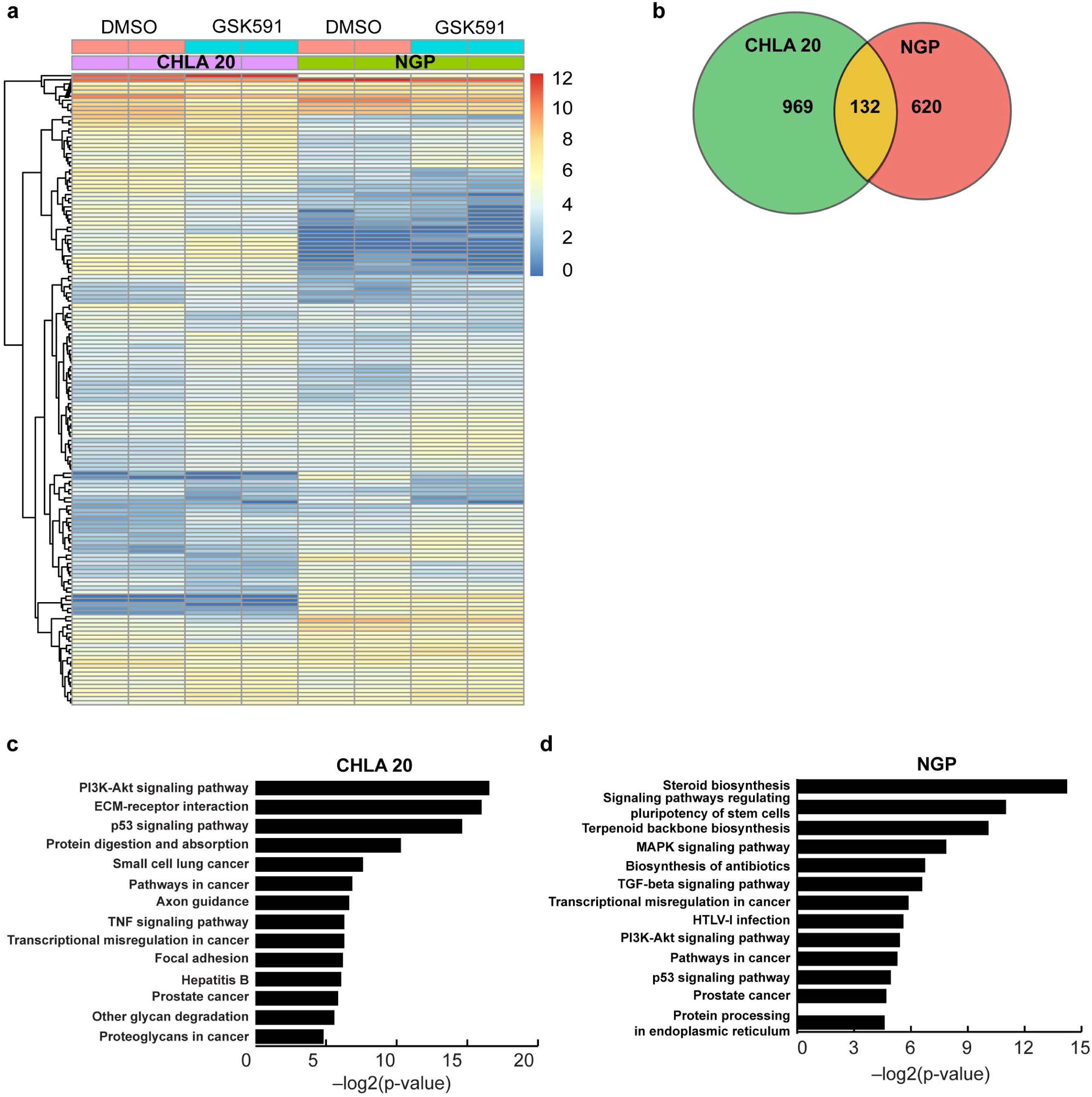
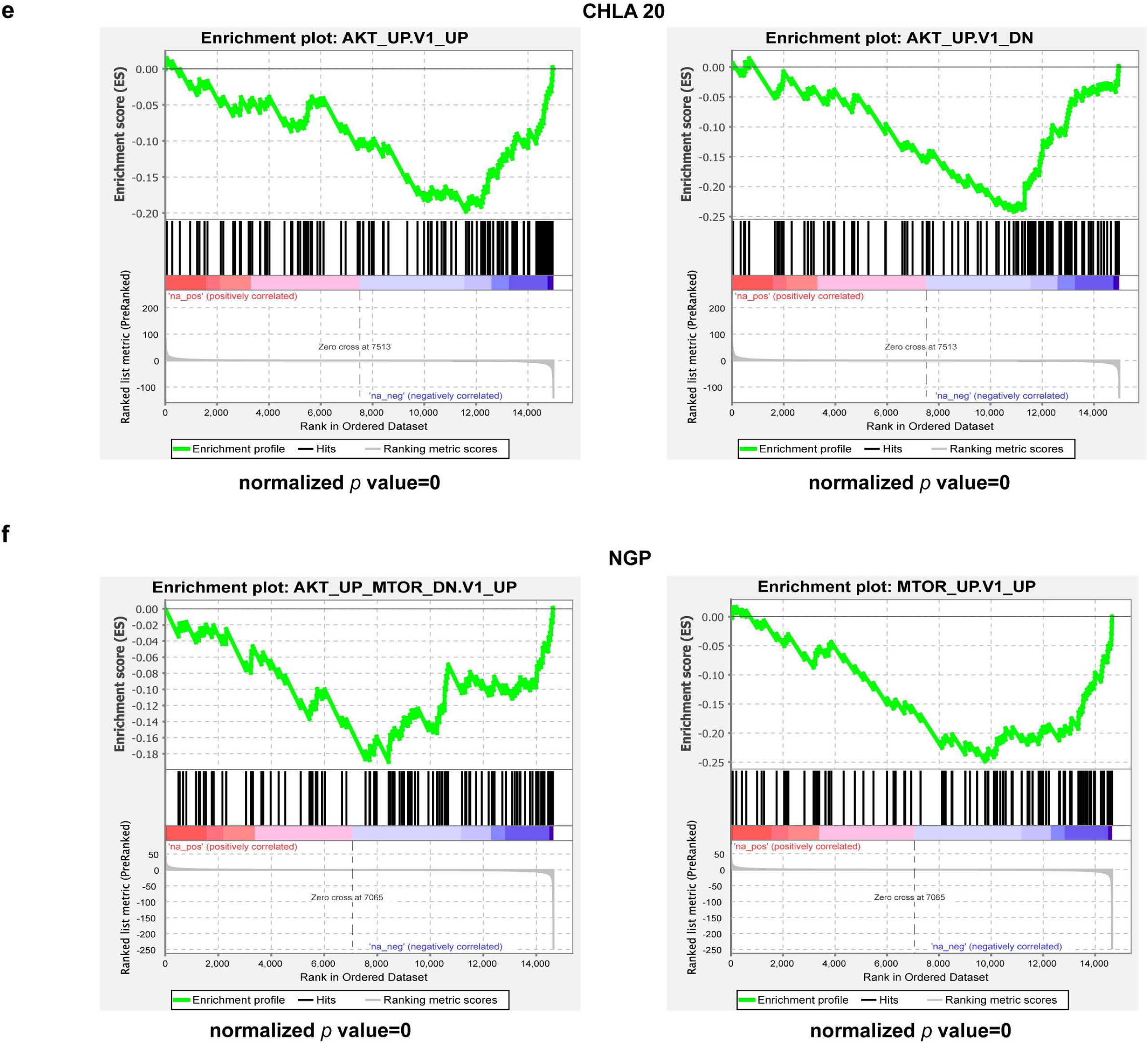
Global transcriptome analysis of PRMT5 inhibition in neuroblastoma cells. **a**, Heat map of significantly differentially expressed genes (fold change >2 and FDR<0.01) between cells treated with DMSO or GSK591. The color scale represented log2 normalized gene expression in DMSO and GSK591 treatment (n = 2 biologically independent samples). **b**, Venn diagram of genes differentially expressed under GSK591 treatment in CHLA20 and NGP (*p*=7.736×10^−48^, hypergeometric test). Enriched Kyoto Encyclopedia of Genes and Genomes (KEGG) pathways analysis in CHLA 20 (**c**) and NGP under GSK591 treatment (**d**). Gene set enrichment analysis of AKT signaling signature genes in CHLA20 cells treated with GSK591 (**e**) and AKT-mTOR signaling signature genes in NGP cells treated with GSK591 (**f**).

### PRMT5 transcriptionally regulates EGFR signaling

Receptor tyrosine kinases (RTKs) regulate pivotal signaling pathways in cancer, such as SRC proto-oncogene, epidermal growth factor receptor (EGFR), fibroblast growth factor receptor (FGFR), and human epidermal growth factor receptor 2 (HER2)^42, 43^. EGFR, together with fibroblast growth factor receptor 4 (FGFR4) and neural growth factor receptor (NGFR), were significantly downregulated under GSK591 treatment, so we validated the expression of these genes by RT-qPCR. As shown in Fig. 4a, the transcripts of EGFR, FRGR4, and NGFR were markedly decreased in GSK591 treated cells. Consistently, the levels of protein encoded by these genes were also significantly downregulated upon inhibition of PRMT5 (Fig. 4b). Furthermore, the expression of these genes was markedly decreased when PRMT5 was depleted (Fig. 4c). EGFR was overexpressed in neuroblastoma patients and associated with resistance to ALK inhibitors and chemotherapy^44–46^. However, the role of EGFR in neuroblastoma has not been as thoroughly investigated as it has in other cancers such as breast cancer, lung cancer, and colorectal cancer^47^.

**Figure 4.**
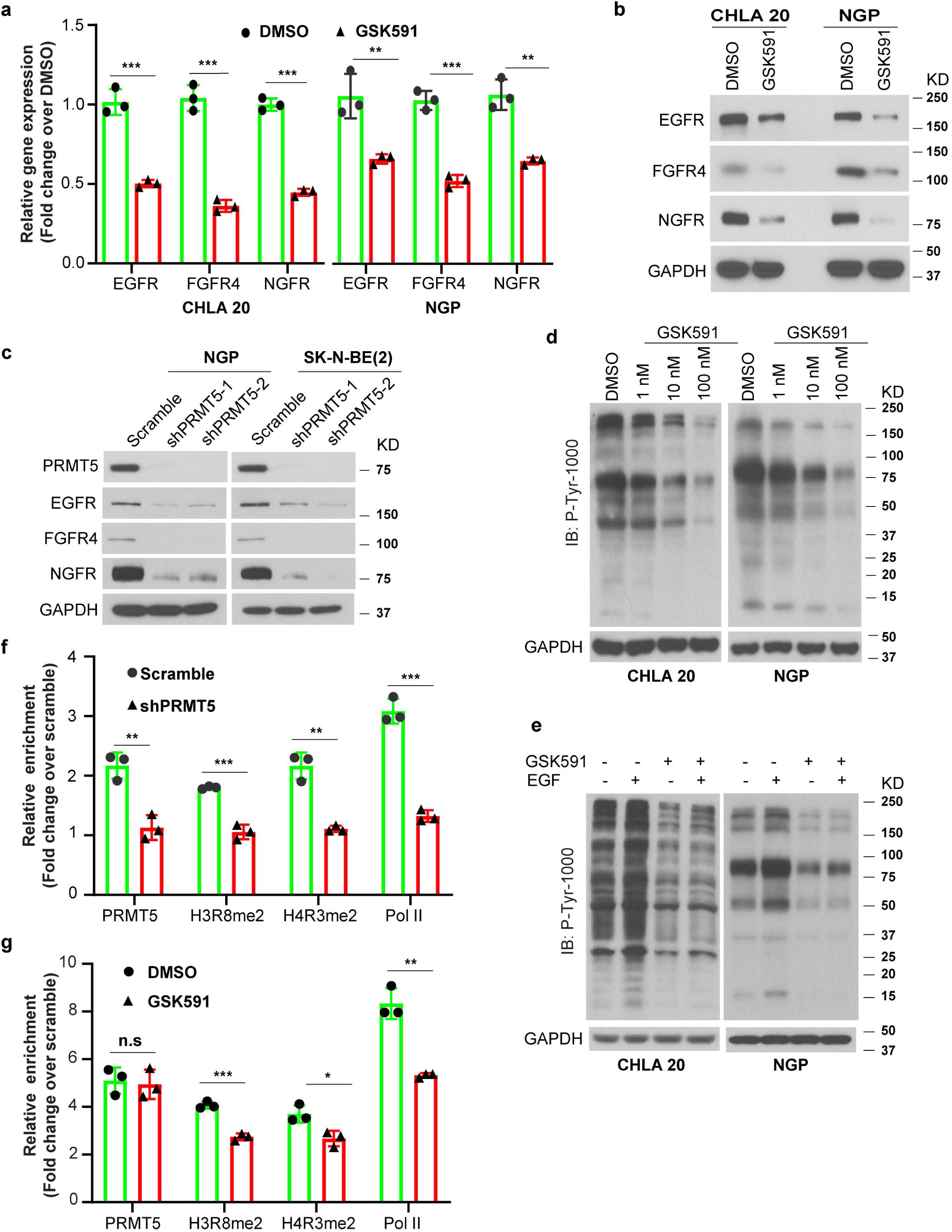
PRMT5 transcriptionally regulates EGFR Signaling. Neuroblastoma cells were treated with DMSO or 100 nM GSK591 for six days. **a**, RT-qPCR analysis of EGFR, FGFR4, and NGFR expression in CHLA20 and NGP cells treated with DMSO or GSK591. **b**, Protein levels of EGFR, FGFR4, and NGFR in neuroblastoma cells treated with DMSO or GSK591. **c**, Western blots of EGFR, FGFR4, and NGFR in control and PRMT5 knockdown cells. **d**, EGFR signaling examined by immunoblotting with Phospho-Tyrosine (P-Tyr-1000) MultiMab™ Rabbit mAb mix in cells treated with DMSO or increasing doses of GSK591. GAPDH was used as a loading control. **e**, The response to EGF stimulation measured by P-Tyr-1000 on cells treated with DMSO or GSK591. The occupancy of PRMT5, H3R8mes2, H4R3mes2, and Pol II at EGFR promoter measured by chromatin immunoprecipitation (ChIP) in NGP cells harboring scramble or shPRMT5 (**f**), and CHLA20 cells treated with DMSO or GSK591 (**g**). **a**-**g**, n=3. Data were analyzed by multiple *t*-test in GraphPad Prism, \**p* < 0.05; \*\**p* < 0.01; *** *p* < 0.001.

We examined EGFR signaling in cells treated with increasing doses of GSK591. As shown in Fig. 4d, global phospho-tyrosine modifications were drastically decreased by the compound in a dose-dependent manner. Moreover, GSK591 treated cells failed to respond to EGF stimulation, suggesting PRMT5 inhibition resulted in impaired EGFR signaling (Fig. 4e). We also tested a group of well-known RTKs, including IGF1R, VEGFR, and ERBB3 on either protein expression or phosphorylation, but none of them were affected by GSK591 treatment (Supplemental Fig. 3a). PRMT5 methylates arginine residues on Histones H3 and H4 with effects on transcription^48–50^. Chromatin immunoprecipitation (ChIP) identified enrichment of PRMT5 at the EGFR promoter. However, this association was diminished in PRMT5 knockdown cells with a concomitant decrease of H3R8 and H4R3 symmetric di-methylarginine and Pol II occupancy (Fig. 4f). Whereas GSK591 treatment did not affect the association of PRMT5 with the EGFR promoter, it did reduce SDMA on H3R8 and H4R3, as well as Pol II binding at this locus (Fig. 4g). Collectively, these results indicate that PRMT5 enzymatic activity is responsible for the epigenetic activation of EGFR expression.

### PRMT5 inhibition impairs AKT signaling

PRMT5 post-translationally modifies proteins, hence modulating their biological functions^34–36, 51^. We explored the consequences of PRMT5 inhibition on global signaling networks by a proteomics-based targeted screen in neuroblastoma cells treated with DMSO or GSK591. The heatmap presented in Fig. 5a shows the expression, modification, and cleavage of crucial proteins involved in 20 signal transduction pathways. The screening confirmed several results shown above (Fig. 1g), such as the increase in cleaved caspase 3, and revealed additional changes including increased phospho-H2AX Ser139 and cleaved Parp, a hallmark of DNA damage and apoptosis, dramatically increased levels of p27, which is a cell cycle inhibitor, and decreased phospho-EGFR after GSK591 treatment (Fig. 4b). Most strikingly, the single most affected target was phospho-AKT Ser473 in both CHLA20 and NGP cell lines (Fig. 5a). When treated with increasing doses of GSK591, both phospho-AKT Thr308 and Ser473 were significantly diminished in a dose-dependent manner (Fig. 5b). This phenomenon was reproduced in PRMT5 knockdown cells, in which AKT phosphorylation at Thr308 and Ser473 was impaired (Fig. 5c). Most importantly, we were able to verify that AKT phosphorylation was attenuated in tumors from GSK595 treated mice in both xenograft models (Fig.5d).

**Figure 5.**
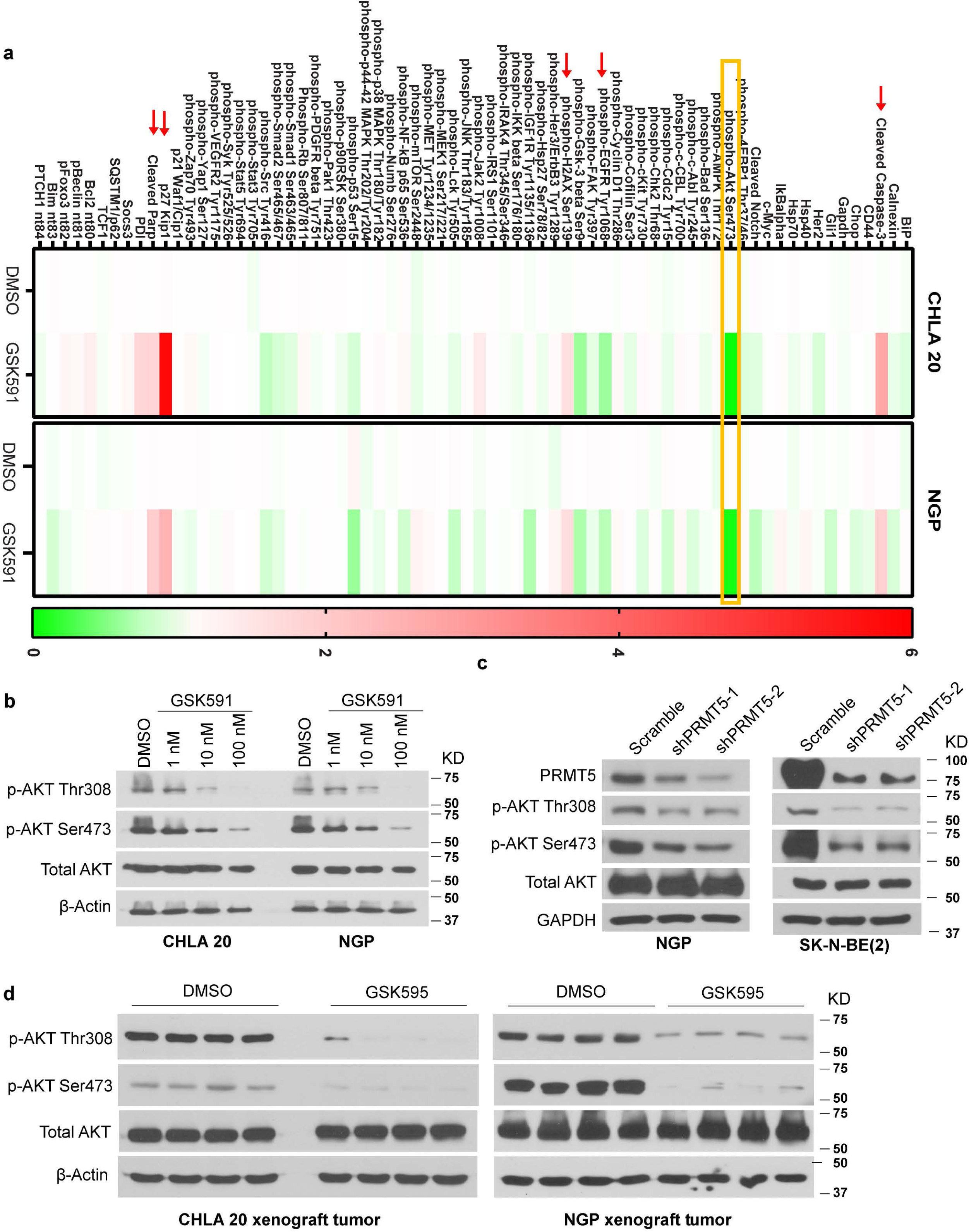
PRMT5 inhibition impaired AKT signaling. **a**, Proteomics-based pathway screening by IPAD platform (Immuno-Paired-Antibody Detection assay) for the expression or modification of key proteins involved in more than 20 signaling pathways. Signals were normalized to internal levels of GAPDH and *beta* Tubulin. Heatmap showed differences between DMSO and GSK591 group based on the average value from two independent experiments (*n* = 2 biologically independent samples). The color scale represented fold changes over DMSO treatment. Representative immunoblots showing phosphorylation of AKT in CHLA20 and NGP cells treated with DMSO or increasing doses of GSK591 (**b**, *n* = 3), in NGP or BE2 cells treated with scramble or shPRMT5 (**c**, *n* = 3), and in CHLA20 and NGP xenograft tumors from vehicle or GSK595 treated mice (**d**, *n* = 4).

### PRMT5 methylates AKT1

Aberrant activation of AKT is driven by oncogenic activation of RTKs, RAS, and PI3K, or by losing tumor suppressor phosphatase and tensin homologue deleted on chromosome ten (PTEN) in many types of cancer^52, 53^. The activity of AKT depends on the dynamic balance between phosphorylation by kinases, such as phosphoinositide-dependent protein kinase 1 (PDK1), rapamycin-insensitive companion of mTOR (Rictor), DNA-dependent protein kinase (DNA-PK), and dephosphorylation by phosphatases, such as protein phosphatase type 2A (PP2A) and PH domain and Leucine-rich repeat protein phosphatases (PHLPP1/2). Besides, PI3K and PTEN modulate the amount of phosphatidylinositol (3,4,5)-trisphosphate (PI(3,4,5)P_3_) thereby, affect AKT activation^54–60^. We hypothesized that PRMT5 inhibition might indirectly regulate AKT activation via these known upstream regulators. However, we failed to detect any changes in these proteins upon GSK591 treatment in neuroblastoma cells (Supplemental Fig. 3b, Supplemental Fig. 4).

Next, we tested the possibility that PRMT5 could directly methylate AKT. In this regard, we detected an interaction between PRMT5 and AKT by co-immunoprecipitation (co-IP) (Fig. 6a). Moreover, PRMT5 co-localized with p-AKT Thr308 and p-AKT Ser473 in control cells, but the co-localization was diminished in GSK591 treated cells. In contrast, the interaction of PRMT5 with total AKT remained unchanged, serving as a matched control for this experiment (Fig. 6b, Supplemental Fig. 5a, b). AKT has three isoforms (AKT1, AKT2, and AKT3) encoded by different genes that share a conserved N-terminal domain, a central kinase domain, and a C-terminal regulatory domain^61^. AKT1 is ubiquitously expressed, AKT 2 is primarily expressed in insulin-responsive tissues, while AKT3 is highly expressed in brain and testes^61, 62^. We examined the abundance of AKT isoforms in neuroblastoma cells and found they were all highly expressed (Supplemental Fig. 3c). In GSK591 treated neuroblastoma cells, symmetric di-methylarginine (SDMA) was markedly decreased on AKT1, modestly reduced on AKT3, but unaffected on AKT2 when pulled down with isoform-specific antibodies (Fig. 6c, Supplemental Fig. 3d). Next, we asked whether the methylation of AKT by PRMT5 correlated with its phosphorylation. After pulldown of endogenous AKT isoforms from cells either treated with DMSO or GSK591, the immunoprecipitated proteins were probed with antibodies recognizing phosphorylation sites on specific AKT isoforms. We showed that indeed phosphorylation of AKT1 at Thr308 and Ser473 was diminished under GSK591 treatment, whereas AKT2 phosphorylation at Ser 474 was unaffected (Fig. 6d, e, Supplemental Fig. 3e, upper and middle panels). Since there are no commercially available antibodies to test the specific phosphorylation sites on AKT3, we probed the immunoprecipitated AKT3 with an antibody recognizing pan-phospho-Serine/Threonine. We did not detect significant changes in AKT3 phosphorylation upon GSK591 treatment (Fig.6d, e, Supplemental Fig. 3e, bottom panels). These results suggest that AKT1 is the primary substrate of PRMT5, and PRMT5 may directly modulate AKT1 activation via methylation.

**Figure 6.**
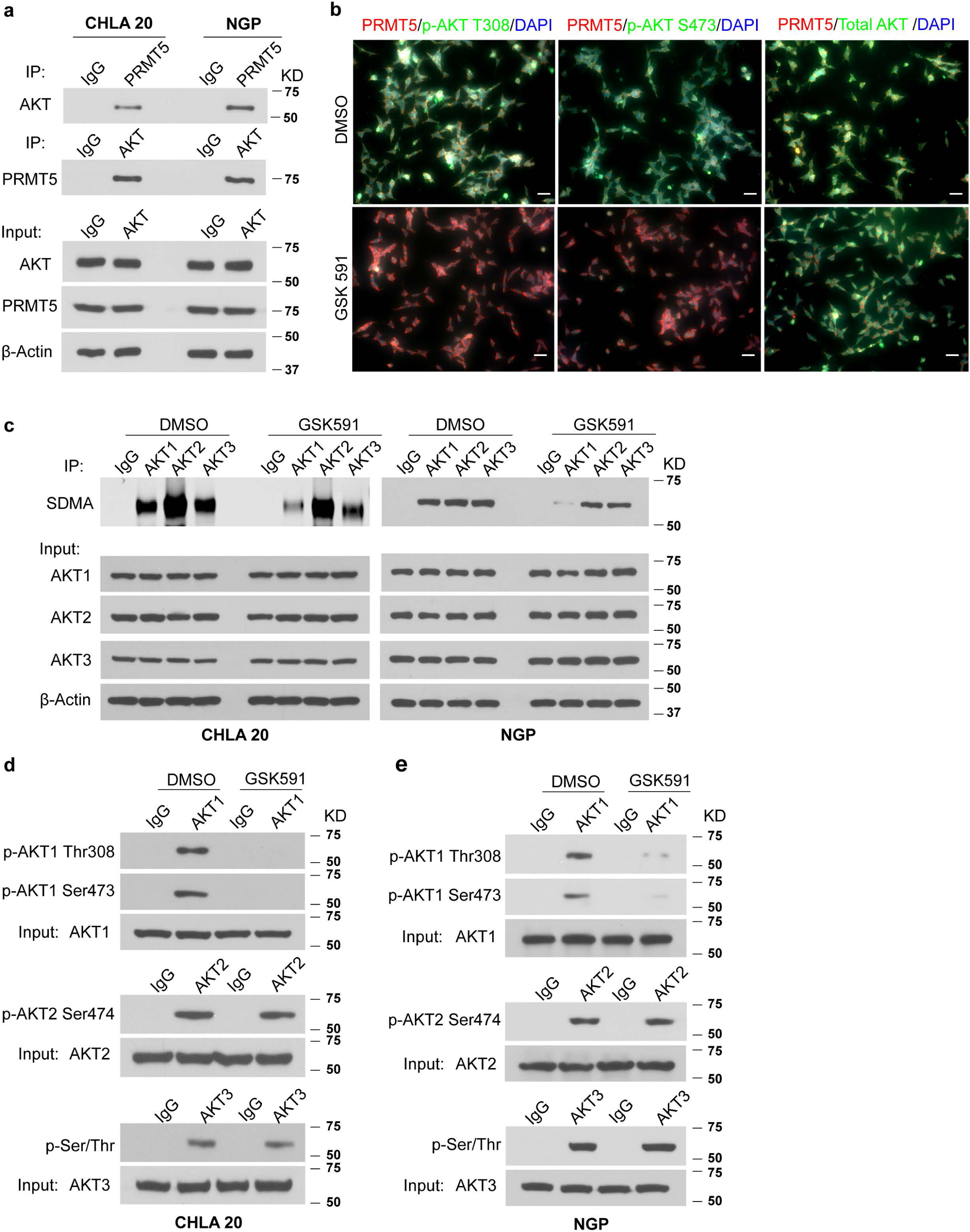
PRMT5 methylated AKT. **a**, PRMT5/AKT interaction captured by co-immunoprecipitation (co-IP). The lysate was IP with anti-PRMT5 antibody followed by immunoblotting with anti-AKT antibody in CHLA20 and NGP cells (top), and the reciprocal co-IP shown in the middle. The input was used as internal control (bottom) (*n* = 3). **b**, Co-localization of PRMT5 with p-AKT Thr308, p-AKT Ser473, and total AKT respectively by immunofluorescence in CHLA20 cells treated with DMSO or 100 nM GSK591 (*n* = 2). Scale bars, 100 *μ*m. **c**, Symmetric dimethylarginine (SDMA) of AKT isoforms in CHLA20 (left) and NGP (right) upon GSK591 treatment examined by immunoprecipitating endogenous AKT isoforms using isoform-specific antibodies followed by immunoblotting with SDMA antibody (*n* = 3). After immunoprecipitated with isoform-specific antibodies, the phosphorylation of AKT isoforms under GSK591 treatment was probed by antibodies specifically recognizing p-AKT1 Thr308 and Ser473 (top), p-AKT2 Ser474 antibody (middle), whereas the phosphorylation of AKT3 was evaluated by a pan antibody recognizing phospho-threonine and serine (bottom) in CHLA20 cells (**d**, *n* = 2) and NGP cells (**e**, *n* = 2) respectively.

### Methylation of AKT1 arginine 15 by PRMT5 is required for AKT1 phosphorylation

Liu *et al*. reported that PRMT5-induced methylation prevented GSK3β-mediated phosphorylation of SREBP1a on S430, leading to its degradation through the ubiquitin-proteasome pathway^51^. In our study, PRMT5 inhibition did not affect the total AKT protein levels (Fig. 5b-d), thereby excluding the possibility that protein degradation reduced AKT phosphorylation. Hence, the decrease of phospho-AKT may be a direct consequence of hypo-methylation under PRMT5 inhibition, which impairs subsequent phosphorylation. We showed above (Fig. 6a) that PRMT5 is associated with total AKT by co-IP, and further confirmed that PRMT5 physically interacted with AKT1 (Supplemental Fig. 3f). PRMT5 preferentially methylates arginine residues within arginine- and glycine-rich (RGG/RG) motifs^36^. Scanning the AKT1 sequence, we identified Arg 15 as a potential site and mutated this arginine to lysine by site-directed mutagenesis. Wild type or R15K-mutant HA-tagged AKT1 was transfected into neuroblastoma cells. After immunoprecipitation with anti-HA antibody-conjugated beads, the methylation and phosphorylation status of wild type or mutant AKT1 were examined. SDMA of mutant AKT1 was abolished compared to its wild type counterpart, indicating Arg 15 is the predominant methylation site of PRMT5 (Fig. 7a, top). Since there was a trace amount of SDMA detected on mutant AKT1, we cannot rule out a minor role of PRMT5 methylating AKT1 on arginine residues other than R15, other type II PRMTs methylating AKT1, or a little background. Notably, phosphorylation of Thr308 and Ser473 on mutant AKT1 was diminished (Fig. 7a, middle). Moreover, expression of wild type but not a catalytically inactive mutant form of PRMT5 could ultimately rescue the decreased AKT activation and EGFR expression in cells treated with GSK591. This result specifies the significance of the methyltransferase activity of PRMT5 in regulating AKT phosphorylation and EGFR expression (Fig. 7b). AKT activation involves two sequential phosphorylation events, including a priming step of phosphorylation on Thr 308 within the activation loop and a full activation step of phosphorylation on Ser 473 in the C-terminal hydrophobic motif. In the R15K mutant AKT1, both phosphorylation events were severely compromised. Our results suggest that methylation of R15 mediated by PRMT5 is required for AKT1 activation.

**Figure 7.**
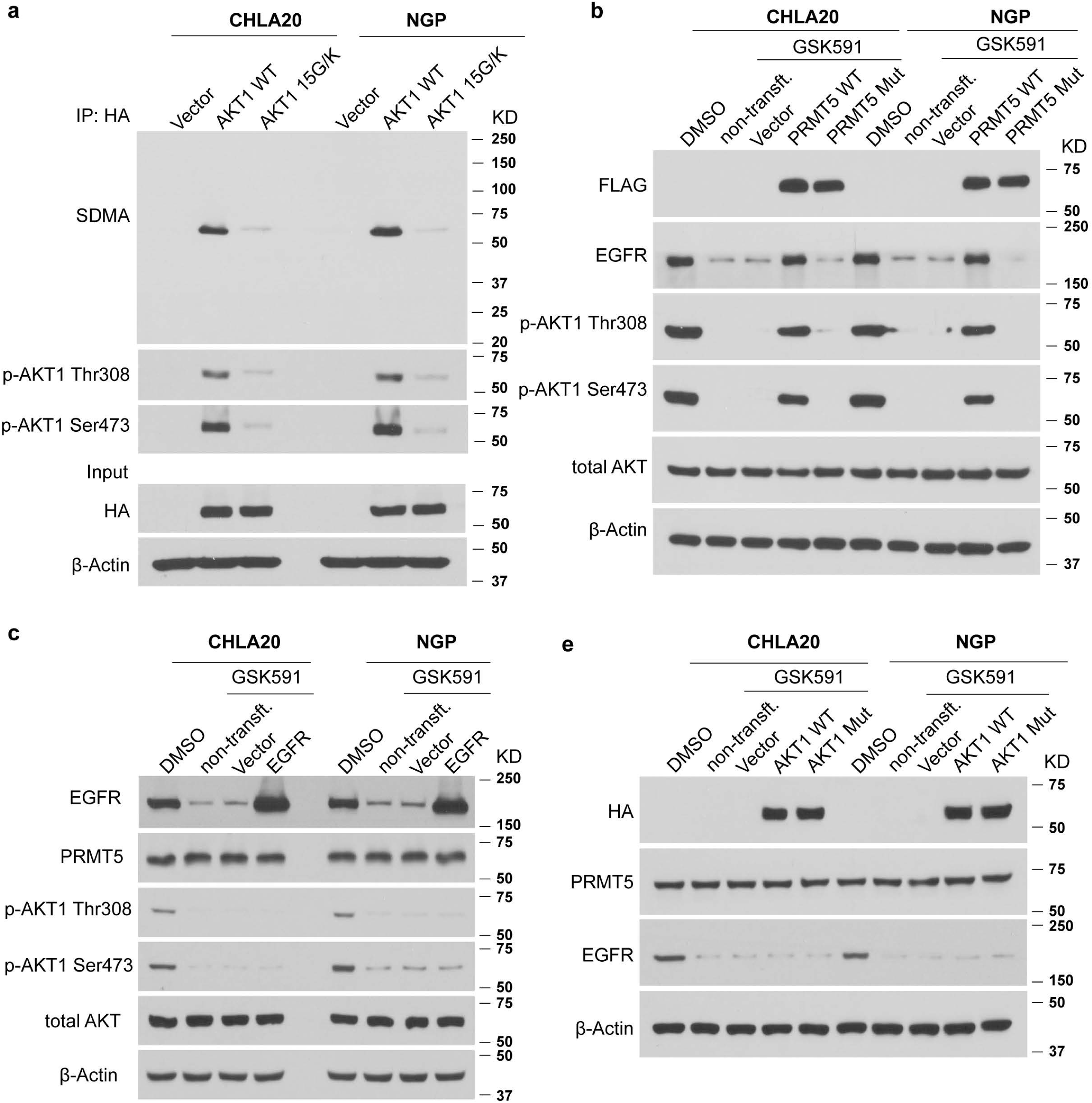
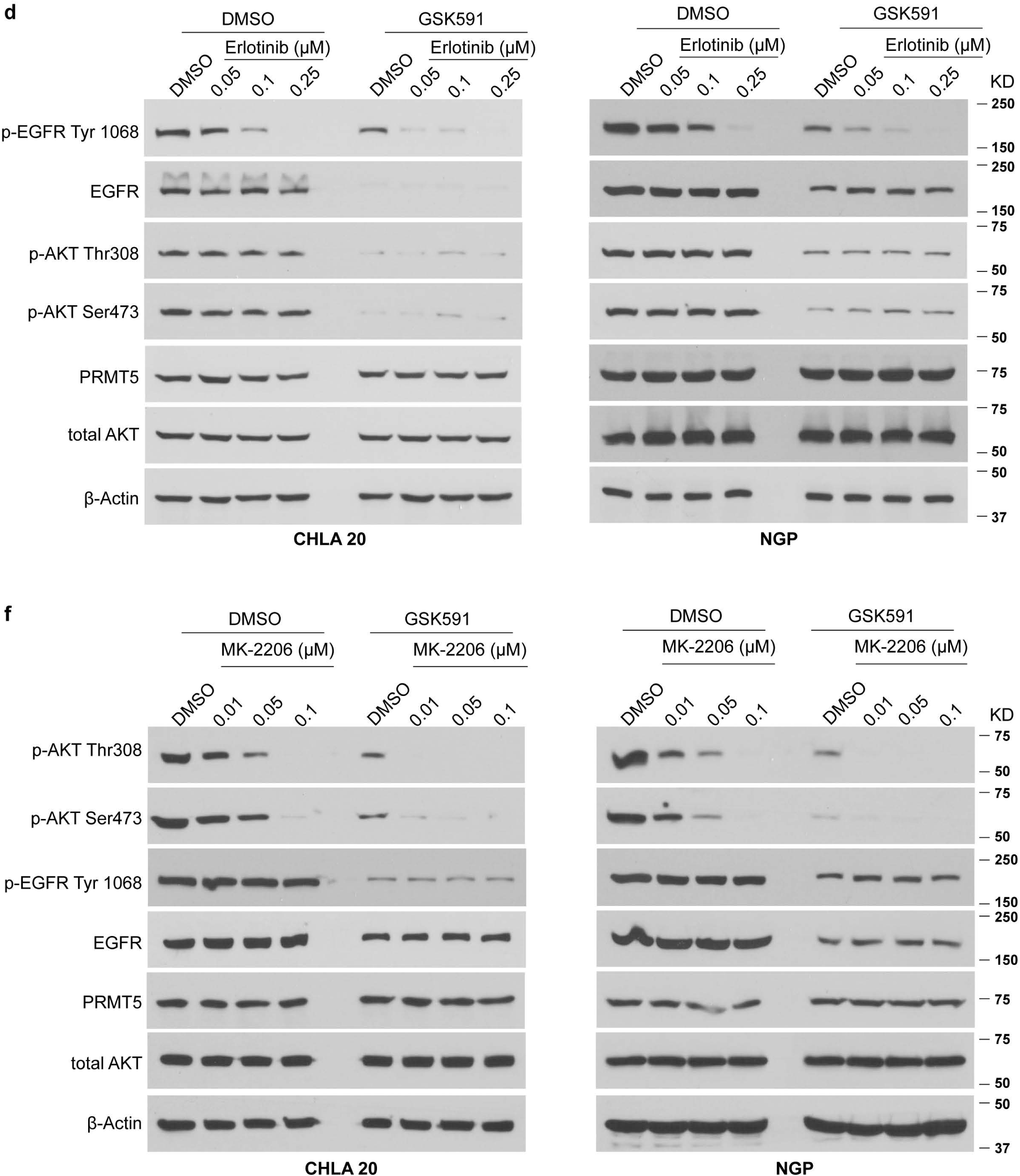
PRMT5 mediated methylation was required for AKT phosphorylation. **a**, Neuroblastoma cells were transfected with vector, wild type or R15K mutant AKT1 and harvested 24 hours post transfection. Exogenous wild type or mutant AKT1 was pulldown by anti-HA antibody conjugated beads. The immunoprecipitated proteins were probed with anti-SDMA, anti-AKT1 phosphorylation antibodies. **b**, **c**, **e**: Neuroblastoma cells were pre-treated with DMSO or 100 nM GSK591 for six days before plated at an equal number of cells for each transfection experiment. **b**, Neuroblastoma cells treated with DMSO or GSK591 were either non-transfected or transfected with vector, wild type or catalytically inactive PRMT5, respectively. Immunoblots showing the levels of exogenous wild type or mutant PRMT5, EGFR, and phosphorylation of AKT1. **c**, Neuroblastoma cells treated with DMSO or GSK591 were either non-transfected or transfected with vector or EGFR, respectively. The expression of EGFR, PRMT5, and phosphorylation of AKT1 examined by Western blot. **d**, **f**: Neuroblastoma cells were pre-treated with DMSO or 100 nM GSK591 for four days before plated at an equal number of cells for subsequent experiments. **d**, Neuroblastoma cells were pre-treated with DMSO or GSK591 and continued with either DMSO or Erlotinib at increasing doses in combination. Immunoblots showing the levels of p-EGFR Tyr 1068, EGFR, p-AKT Thr308, p-AKT Ser473, and PRMT5. **e**, Neuroblastoma cells pretreated with DMSO or GSK591 were either non-transfected or transfected with vector, wild type or methylation mutant AKT1. The protein levels of exogenous wild type or mutant AKT1, endogenous PRMT5 and EGFR analyzed by Western blot. **f**, Neuroblastoma cells were pre-treated with DMSO or GSK591 for four days and continued with either DMSO or MK-2206 at increasing doses in combination. The levels of p-AKT Thr308, p-AKT Ser473, p-EGFR Tyr 1068, EGFR and PRMT5 examined by immunoblotting. **a**-**f**, *n* = 3.

### PRMT5 directly and independently regulates EGFR and AKT activation

Receptor tyrosine kinases are critical inducers of AKT signaling, as activation of PI3K signaling leads to AKT activation^26^. Since we showed that EGFR signaling was attenuated in neuroblastoma cells under PRMT5 inhibition, we next questioned whether this could contribute, at least partially, to the diminished AKT activation. Introducing exogenous EGFR in GSK591 treated cells did not restore the decreased phosphorylation of AKT at Thr308 and Ser473, suggesting that EGFR does not directly contribute to the deficiency of AKT activation (Fig. 7c). To further explore the impact of endogenous EGFR signaling on AKT activation, we blocked EGFR function with the EGFR inhibitor erlotinib alone or in combination with GSK591. Erlotinib treatment attenuated EGFR activation in a dose-dependent manner, as evidenced by the decreased EGFR autophosphorylation at Tyr 1068 (Fig.7d, top panel). However, erlotinib failed to reduce AKT phosphorylation, nor did it affect protein levels of PRMT5 or EGFR (Fig.7d, middle). These results suggest that AKT activation is not downstream of EGFR in the conditions we tested. AKT has been shown to facilitate the trafficking and recycling of EGFR^63^. To determine whether the impaired AKT activation by GSK591 influences EGFR expression, we transfected GSK591 treated cells with wild type or R15K mutant AKT1. Neither the wild type nor the mutant AKT1 affected the protein level or the phosphorylation of EGFR (Fig. 7e). To further delineate the regulatory circuit of PRMT5, EGFR, and AKT, we treated neuroblastoma cells with AKT inhibitor MK-2206 alone or in combination with GSK591. MK-2206 treatment significantly blocked AKT phosphorylation, indicating the effectiveness of MK-2206 under the doses used (Fig. 7f). However, the addition of MK-2206 failed to affect protein levels of EGFR or PRMT5, nor the phosphorylation of EGFR (Fig. 7f). Our results indicate that PRMT5 acts directly to regulate EGFR transcriptionally and post-translationally modulate AKT activation by distinct mechanisms.

### PRMT5 regulates EMT networks through EGFR and AKT

Previous studies have shown that AKT acts as a mediator downstream of TNFα and TGFβ in regulating the transcriptional activity of EMT-TFs such as SNAIL and TWIST1^64–66^. EGFR signaling is one of the critical transduction mechanisms underlying the induction of SNAIL-dependent EMT^67–69^. Given that EGFR signaling and AKT activation are compromised under PRMT5 inhibition, we speculated that the EMT program is likely to be attenuated. Indeed, the expression of EMT-TFs such as ZEB, TWIST1, and SNAIL declined in cells treated with GSK591 in a dose-dependent manner (Fig. 8a). Moreover, these protein levels of these EMT transcription factors were reduced when PRMT5 was knocked down (Fig. 8b). Importantly, diminished levels of ZEB, TWIST1, and SNAIL were detected in xenograft tumors from GSK595 treated mice compared to control mice (Fig. 8c).

**Figure 8.**
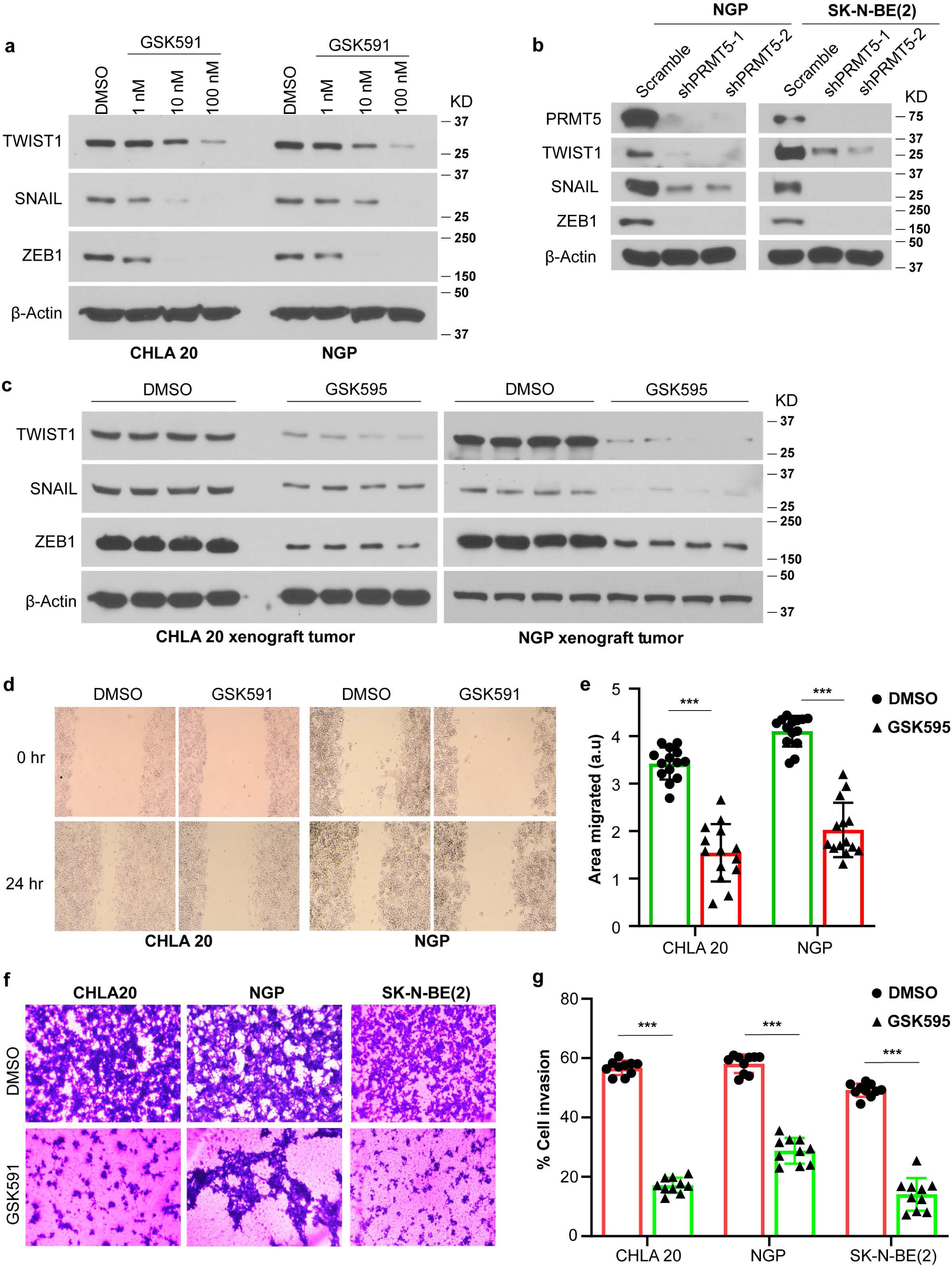
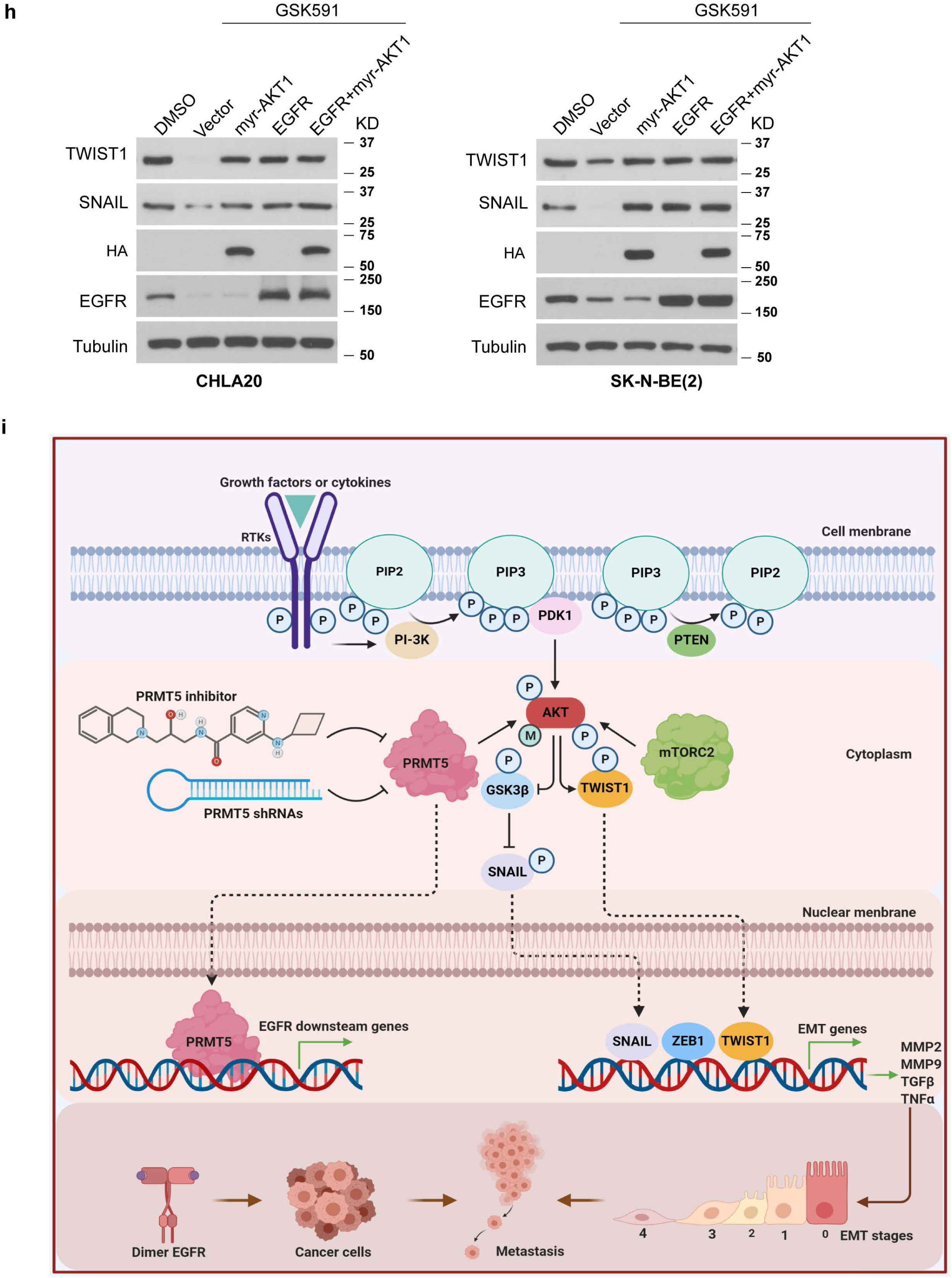
PRMT5 regulates EMT networks through EGFR and AKT. Immunoblots showing the expression of TWIST1, SNAIL, and ZEB1 in neuroblastoma cells treated with DMSO or increasing doses of GSK591 (**a**, *n* = 3), in xenograft tumors from mice treated with vehicle or GSK595 (**b**, *n* = 4), in the scramble and shPRMT5 cells (**c**, *n* = 3). **d**, Representative images of cells migrated to the cleared space (wound) after 24 hours. **e**, Quantification of *in vitro* cell migration assay. Five images per sample were analyzed. The distance between the edges of the wound was measured at time 0 and 24 hours, and the reported migrated distance corresponding to the difference between 0 and 24 hours. The migration area was determined by measuring the total area of the wound using the ImageJ software. **f**, Representative images of cells invaded to ECM coated membrane in trans-well invasion assay. **g**, Percentage of invasive cells normalized by cell numbers in the non-ECM coated 12-well plate using Image J. **h**, Neuroblastoma cells were pre-treated with DMSO or 100 nM GSK591 for six days and plated at an equal number of cells for each transfection experiment. Cells were transfected with vector, constitutively activated AKT1 (myr-AKT1), EGFR alone or in combination, respectively. Protein levels of TWIST1 and SNAIL examined by immunoblotting (n=3). **i**, Graphical summary. **e** & **g**, statistics were performed using unpaired *t*-test in GraphPad Prism.

Moreover, *in vitro* cell migration and invasion assays confirmed the biological consequences of the decreased EMT. Cell migration was attenuated in GSK591 treated cells compared to control cells, where the impact from cell proliferation was eliminated by mitomycin C treatment (Fig. 8d, e). Most importantly, cell invasion through the extracellular matrix was significantly reduced under GSK591 treatment (Fig. 8f, g). These results are consistent with our *in vivo* study, where PRMT5 inhibition blocked tumor cell metastasis to the liver.

If PRMT5 regulates EMT via EGFR and AKT signaling pathways, redeem the levels of EGFR or/ and AKT activation should restore the EMT program. We transfected GSK591-treated cells with either vector, EGFR, constitutively activated AKT1 (myr-AKT1) alone or in combination. And then, we analyzed the expression of EMT TFs. As shown in Fig. 8h, overexpressing EGFR, or myr-AKT1 alone is sufficient to rescue the decreased expression of TWIST and SNAIL. This result indicates that both EGFR and AKT1 signaling pathways regulate the expression of EMT-TFs, and they are capable of compensation for the loss of either one. Our results suggest that PRMT5 promotes neuroblastoma metastasis by increasing the transcription of EGFR and elevating AKT1 activation by methylation on arginine 15 (Fig. 8i).

## Discussion

Metastatic neuroblastoma represents a major clinical challenge, as less than 50% of children with high-risk aggressive cancer will survive^24^. While intensive chemotherapy regimens and immunotherapeutics improve cure rates, this comes with increased short- and long-term toxicity. Relapse of drug-resistant, metastatic disease remains the primary cause of death for these very young children. Thus, novel approaches to prevent and treat metastatic disease are urgently needed.

PRMT5 is the primary symmetric dimethylarginine methyltransferase. PRMT5 overexpression has been observed in a number of highly aggressive metastatic cancer types^35, 40^. We illustrate here that PRMT5 inhibition attenuates primary tumor growth and blocks hepatic metastasis (Fig. 2). We show that PRMT5 transcriptionally regulates EGFR (Fig. 4) and modulates AKT1 activation via direct methylation on Arg 15 (Fig. 5, 6, 7), both fundamental components of critical signaling networks that orchestrate the EMT program (Fig. 8). In line with the inhibition of liver metastasis in xenograft models, we confirm that PRMT5 inhibition decreases the expression of EMT-TFs and impairs cell migration and invasion (Fig. 8). Our findings collectively prompt us to propose PRMT5 as a very promising and druggable target for not only neuroblastoma treatment but also other high-risk tumors, with higher potential to develop metastatic tumors.

We showed that PRMT5 is associated with EGFR promoter and methylates H3R8 and H4R3, consequently regulating EGFR expression (Fig. 4). High levels of EGFR are associated with poorer prognosis and outcomes in diverse tumor types, and therapies targeting the kinase activity of EGFR have been vigorously tested in clinical trials^70^. However, rapidly acquired mutations compromise the efficacy of these drugs^71^. Our results suggest that inhibiting PRMT5 to reduce the protein levels of EGFR represents a complementary/alternative approach to block EGFR signaling. Indeed, the combination of PRMT5 inhibitor GSK591 and EGFR inhibitor erlotinib abolished EGFR autophosphorylation (Fig. 7). Hsu *et al*. reported that PRMT5 methylated EGFR Arginine 1175, and this modification positively modulated EGF-induced EGFR trans-autophosphorylation at Tyr 1173 in breast cancer cell lines^72^. But the depletion of PRMT5 by siRNA did not downregulate EGFR protein levels, nor did it affect AKT phosphorylation^72^. These results are in contrast to our observations, which may suggest context-dependent or tissue-specific effects and functions for PRMT5.

Previous studies have linked PRMT5 to EMT program^73–75^. Chen *et al*. reported that PRMT5-MEP50 activity is required for TGFβ induced EMT by direct association with and methylation of histone substrates at EMT gene loci^18^. In the RNA-seq datasets, we detected a modest but significant decrease of ZEB1, SNAIL, and TWIST1 in GSK591 treated CHLA20 cells, whereas only SNAIL transcripts were downregulated in GSK591 treated NGP cells. Nevertheless, all protein levels of these EMT-TFs were markedly reduced in both cell lines, suggesting that the compromised expression of EMT-TFs is primarily regulated at the posttranslational rather than the transcriptional level. Although PRMT5 inhibition decreased AKT phosphorylation, we did not detect a significant change in mTOR protein level or reduced phosphorylation of Ser2448, a direct target of AKT^76^ (data not shown). Overall, our results suggested a direct role of AKT as a modulator of the EMT program that is independent of mTOR.

Our study revealed that PRMT5 mediated methylation on AKT1 is required for AKT1 activation (Fig.7). Discovered almost three decades ago, many downstream effectors of AKT have been characterized, such as GSK3, mTOR, and FOXO^76, 77^. In contrast, only a few upstream regulators have been confirmed, including PIP3 modulators PI3K and PTEN^54, 57, 78^, kinases PDK1, Rictor, and DNA-PK^54, 56, 59, 79^, and phosphatases PP2A phosphatases and PHLPP^58, 80^. Despite both *in vitro* and *in vivo* results showing that PRMT5 inhibition markedly decreased AKT phosphorylation at Thr 308 and Ser 473, none of the previously defined AKT activators were altered by the treatment (Fig. 5, Supplemental Fig. 2). Although we can’t completely rule out the existence of an unknown upstream regulator of AKT being PRMT5’s target, it is unlikely that PRMT5 modulates AKT activation via an indirect mechanism. Nevertheless, we detected the association of PRMT5 with total AKT, p-Thr308 AKT, and p-Ser473 AKT (Fig. 6, Supplemental Fig.5). Further, we showed that reduced symmetric dimethylarginine on endogenous AKT1 correlates with its decreased phosphorylation under PRMT5 inhibition (Fig. 6). Thus we confirmed that AKT1 is the primary target of PRMT5 in the context of neuroblastoma. Notably, phosphorylation on Thr308 and Ser473 was diminished when AKT1 Arg 15 was mutated to Lysine. These results provide substantial evidence that PRMT5 mediated arginine methylation of AKT1 is necessary for its full activation (Fig.7). The development of AKTinhibitor has evolved from targeting ATP-binding to allosteric sites and catalytic activation loop^81^. Our findings may offer a novel strategy to develop a new generation of inhibitory agents to block the oncogenic function of AKT.

We show *in vitro* and *in vivo* that PRMT5 is an upstream regulator of EGFR and AKT signaling pathways, essential for cancer cell proliferation and metastasis. It is noteworthy that overexpressing EGFR or constitutively activated AKT1 alone is sufficient to restore the decrease of EMT-TFs. This finding highlights the significance of PRMT5 inhibition in targeting the EMT program to block metastasis. In combination with PRMT5 inhibition, both Erlotinib and MK-2206 are more effective at lower doses and achieved better inhibition of EGFR and AKT, respectively. Our results provide novel mechanisms linking PRMT5 to EGFR and AKT regulation and merit further exploration of potential therapeutic utility for metastatic cancers.

## Acknowledgments

We thank Drs. Anthony Imbalzano, Jeffrey Nickerson, Ivana de la Serna, Thomas Fazzio, Heidi Tissenbaum, and Michael Green for their invaluable insights to improve this manuscript. This work is supported by Dean’s Research Fund, University of Massachusetts Medical School, and Hyundai Scholar Hope Grant from Hyundai Hope on Wheels Foundation awarded to Dr. Jason Shohet.

## Material and Methods

### Chemicals

PRMT5 methyltransferase activity inhibitor GSK3203591 was purchased from Cayman Chemical. The *in vivo* active sister compound GSK3326595 was purchased from Chemietek. Doxorubicin and etoposide were purchased from Sigma-Aldrich. All chemicals were dissolved in 100% DMSO as stock solution and subsequently diluted in phosphate-buffered saline or growth media to working solution.

### Cell culture

CHLA-20 and SK-N-BE (2) cell lines were obtained from The Children Oncology Group. CHLA-20 cells were cultured in IMEM medium (Invitrogen, San Diego, CA, USA) supplemented with 10% fetal bovine serum (FBS) (Sigma-Aldrich, St. Louis, MO), 100 μg/ml streptomycin (Gibco), and ITS (5 μg/mL insulin, 5 μg/mL transferrin and 5 ng/ml sodium selenite, Invitrogen). SK-N-BE (2) and NGP cells were grown in RPMI1640 medium (Invitrogen) supplemented with 10% fetal bovine serum (FBS) (Sigma-Aldrich), 100 μg/ml streptomycin.

### Plasmids

Human pCDNA3/Flag-HA-AKT1, Plncx/myr-HA-AKT1, pCDNA6A/EGFR, and pHIV-iRFP720-E2A-Luc were purchased from Addgene. pCDNA3.1/Flag-PRMT5 wild type and dead enzyme mutant were as described previously^39^. pCDNA3/Flag-HA-AKT1 15G/K mutant was generated by site mutagenesis using QuickChange Lightning Site-Directed Mutagenesis Kit following the manufacturer’s protocol (Agilent). Primers introducing mutation sites were designed by Agilent PrimerDesign Program on Agilent website listed in Table 3. pLKO-Tet-ON inducible shPRMT5 constructs were generous gifts from Dr. William R. Sellers (Broad Institute).

### Stable cell lines

CHLA20 and NGP cells were transduced with pHIV-iRFP720-E2A-Luc and enriched by FACS sorting on RFP 700 channels. NGP and SK-N-BE (2) cells were transduced with pLKO-Tet-ON inducible scramble or shRNA targeting PRMT5 followed by puromycin selection as previously reported^82^.

### Fluorescent activated cell sorting (FACS)

Primary cells were isolated from xenograft tumors and liver. The percentage of iRFP 720 positive cells was analyzed by FACS. iRFP720 positive cells from the primary tumor were enriched by FACS sorting for transcriptome analysis by RNA-seq.

### Cell viability assay

Cells were seeded in 96-well plates and then maintained in the presence of vehicle or increasing doses of GSK591 for six days before the addition of 20 μL CellTiter 96 AQueous One Solution per well (Promega Corporation, Madison, WI). Plates were incubated for 2 hours before absorbance at OD490 was measured with a Synergy H4 Hybrid microplate reader (Bio Tek, Winooski, VT). Wells containing medium only were used as background for the measurement.

### Hoechst 33342 staining

The cells were treated with DMSO or GSK591 100 nM for six days and then were stained with Hoechst 33342 for 10 min at 37°C. The cells were photographed at 40× magnifications with fluorescence microscopy from Carl Zeiss Microscopy. All images were captured on an AXIO Observer microscope with a ZEISS Axiocam 506 mono camera and Zen 2 Pro software.

### Caspase-3/7 Staining

Apoptosis was measured by caspase-3/7 staining using CellEvent® Caspase-3/7 Green ReadyProbes® Reagent (ThermoFisher Scientific Inc., cat# R37111). CHLA20 and NGP cells were grown until confluence on 6-well plates with six days of treatment with DMSO or GSK591. Cells were fixed in 4% formaldehyde for 20 min and followed by methanol 100% for 10 min. Cells were stained with two drops of Caspase-3/7 green reagent per mL of media for 30 min at room temperature, and the nuclei were stained with DAPI (Sigma-Aldrich, cat#D9542) for 15 min in PBS. The images were taken under Zeiss microscope with a 10× objective. The Caspase-3/7 activity was determined by analyzing particles using the ImageJ software. The percentage of Caspase-3/7 positive cells was normalized to the nuclei stained by DAPI.

### Xenograft mouse model

NOD/SCID mice were used in this study. One million iRFP 720-luciferase transduced CHLA20 or NGP cells suspended in 0.1 ml of PBS were surgically implanted in the left renal capsule of mice. Tumor growth was monitored weekly by bioluminescent imaging (IVIS Lumina XR System, Caliper Life Sciences, Hopkinton, MA, USA). After ten days, mice bearing tumors with similar sizes (determined by live animal imaging of bioluminescence) were randomly divided into either a “vehicle control” group (0.5 methylcellulose, Sigma-Aldrich #M0430) or a PRMT5 inhibitor “GSK595 treatment” group (100 mg/kg). Animals were treated for two weeks, with 100 mg/kg GSK595 or vehicle by oral gavage twice daily. The bodyweight of mice was monitored weekly. At the end of the treatment, all mice were euthanized. Tumors and the right kidneys (control) were dissected and weighed. Tumor/kidney and liver were subjected to organ imaging by bioluminescence. All procedures were approved by the Institutional Animal Care and Utilization Committee (IACUC) at UMass Medical School.

### Western blot

Tissue or cells were washed with cold PBS and lysed in 1% NP40 lysis buffer containing a protease and phosphatase inhibitor cocktail (Roche Diagnostics, Indianapolis, IN, USA). Proteins were analyzed by SDS-PAGE, followed by immunoblotting with primary antibodies. Blots were incubated with horseradish peroxidase-conjugated anti-goat IgG, anti-mouse or anti-rabbit IgG (Santa Cruz Biotechnology) antibodies, and proteins detected by enhanced chemiluminescence (ThermoFisher Scientific). Antibodies used in this study were listed in Table 1.

### RT-qPCR

Total RNA was extracted from cells using Trizol reagent (Invitrogen) and purified by Direct-zol RNA Purification Kit (Zymo Scientific). cDNA was synthesized by SuperScript Reverse Transcriptase III kit (Invitrogen). The cDNA was amplified in 96-well reaction plates with a SYBR green PCR Master Mix (Applied Biosystems) on an ABI 7500 real-time PCR thermocycler. The sequences of forward and reverse primers are listed in Supplementary Table 1. The relative level of target transcripts was calculated from duplicate samples after normalization against human 45S pre-RNA. Dissociation curve analysis was performed after PCR amplification to confirm primers’ specificity. Relative mRNA expression was calculated using the ΔΔC_T_ method.

### Immunofluorescence

Cells were treated with DMSO or GSK591 100nM for six days and fixed with 4% paraformaldehyde for 20 min at room temperature, followed by permeabilization in ice-cold methanol for 10 min at −20°C. Cells were then incubated in blocking buffer (5% BSA in PBS) for 1 h at room temperature, followed by incubation with anti-PRMT5 antibody (Santa Cruz Biotechnology, catalog no. sc-376937) and anti-Thr308-AKT, anti-Ser473-AKT and total-AKT antibody at 4 °C overnight. Cells were washed three times for 5 min in PBS, then incubated for two hours with Alexa Fluor 488-conjugated goat anti-rabbit secondary antibody against phospho-AKT and total-AKT and Alexa Fluor 596-conjugated goat anti-mouse secondary antibody for PRMT5 (diluted 1:100 in blocking buffer, Invitrogen) at room temperature. Finally, the nuclei were stained with DAPI (Sigma, cat#10236276001) for 30 min at room temperature before visualization. Cells were observed with ZEISS Axiocam 506 mono Digital Camera for Fluorescence Microscopy.

### Co-Immunoprecipitation (Co-IP)

CHLA-20 and SK-N-BE (2) cells were seeded in 10 cm culture dishes. Proteins were extracted from those cells using lysis buffer (20 mM Tris, PH 7.4, 150 mM NaCl, 2 mM EDTA, 2 mM EGTA, 1 mM sodium orthovanadate, 50 mM sodium fluoride, 1% Triton X-100, 0.1% SDS and 100 mM phenylmethylsulfonyl fluoride) and centrifuged at 13000 rpm for 10min at 4°C. The cell lysates were pre-cleared with protein A or protein G beads at 4oC for 1 hour. Subsequently, 5 μg primary antibody or isotype IgG was added into the cleared cell extracts and incubated at 4°C for overnight. Protein A or protein G beads were added into the cell extracts and incubated at 4°C for 3 hours. Then the beads were washed three times with wash buffer (100 mM NaCl, 50 mM Tris pH 7.5, 0.1% NP-40, 3% glycerol, 100 mM phenylmethylsulfonyl fluoride). Finally, the beads bound proteins were eluted by adding 2x Laemmli sample buffer and heated at 95C for 5 minutes. Eluted proteins were analyzed by western blotting.

### Bioinformatics analysis

The gene expression and clinical data from 161 neuroblastoma patients were downloaded from the TARGET database (https://ocg.cancer.gov/programs/target/data-matrix). Three sets of microarray data from neuroblastoma patient cohorts were downloaded from R2 database (R2: Genomics Analysis and Visualization Platform(http://r2.amc.nl). The relative expression of PRMT5 and survival analysis was performed on these two groups using the survminer R package.

### RNA-seq

CHLA-20 and NGP cells were treated with vehicle or IC50 dose of GSK591 for six days. Dead cells were removed by Dead Cell Removal Kit (Miltenyi). Total RNA was extracted from live cells using Trizol reagent (Invitrogen) and purified by Direct-zol RNA Purification Kit (Zymo). RNA integrity was assessed using the RNA Nano 6000 Assay Kit of the Bioanalyzer 2100 system (Agilent Technologies, CA, USA). The mRNA enrichment, mRNA fragmentation, second-strand cDNA synthesis, size selection, PCR amplification, and sequencing were performed by BGI using DNBseq sequencing technology (One Broadway, Cambridge, MA 02142). RNA-seq reads from each sample were aligned using STAR against the GRCh38/hg38 human reference genome^83^. DESeq2 was used to identify differentially expressed genes using raw read counts quantified by htseq-count with GENCODE V29 gene annotation. GO term enrichment on differentially expressed genes was performed using DAVID^84–86^. Gene set enrichment analysis (GSEA) for differentially expressed genes was performed using pre-ranked gene lists ordered by −log10 (P value) multiplied by +1 for upregulation or −1 for downregulation^87^.

### ActivSignal IPAD assay

The signal transduction pathway analysis was performed as previously reported^39^. Briefly, the whole-cell lysate was submitted to ActivSignal for analyzing the expression or phosphorylation of 70 pivotal proteins involved in more than 20 signaling pathways. The signal was normalized to the expression of house-keeping genes. Each pathway was covered by multiple targets. Samples were analyzed in independent biological duplicates.

### Cell migration assay

*In vitro* cell migration assay was assessed by wound healing. CHLA20 and NGP cells were grown until confluence in 6 well plates with six days of treatment of DMSO or 100 nM GSK591. Then cells were incubated for two hours with 10 µg/ml mitomycin C (Sigma-Aldrich) to inhibit cell proliferation before the wound generation. Cells were maintained in media containing 10 µg/ml mitomycin C afterward. A sterilized 200 µl pipette tip was used to generate wounds across the cell monolayer, and debris was removed by washing with media. The monolayers were then incubated for 24 h at 37 °C. The progress of cell migration into the wound was monitored using a Zeiss microscope with a 10× objective. The bottom of the plate was marked for reference, and the same field of the monolayers was photographed at 0 and 24 hours. Five images per sample were analyzed. The distance between the edges of the wound was measured at time 0 and 24 h, and the reported migrated distance corresponds to the difference between these two. The migration area was determined by measuring the total area of the wound using the ImageJ software.

### Cell invasion assay

*In vitro* invasion assay was performed in cell culture inserts with 8 µm pore size (BD Biosciences, MA, USA) coated with extracellular matrix (ECM) from Engelbreth-Holm-Swarm murine sarcoma (Millipore Sigma, cat#E1270). The basement membrane of the inserts was coated with a 50 μL of ECM gel at room temperature, then hydrolyzed at 37°C for 30 min. CHLA20, NGP, and SK-N-BE(2) were treated with either DMSO or 100 nM GSK591 for six days, then detached and suspended at 5×10^4^ cells/ml in serum-free media containing DMSO or 100 nM GSK591, respectively. Two hundred microliter cell suspension was added onto the upper surface of the insert. Seven hundred and fifty microliter growth media were added to the bottom chamber. After 15 h incubation, cells grown on the top surface of the insert were removed with cotton swabs. Cells that invaded to the opposite surface of the insert were washed with PBB, fixed in 4% formaldehyde for 5 min and followed by methanol 100% for 20 min, then stained with 0.25 % crystal violet for 10 min. Photographs were taken under a light microscope and quantified using Image J software. The percentage of cells of invasion was normalized to the equal number of cells plated in 24-well plate cultured for the same period.

### Statistical Analyses

All quantitative data points represent the mean of three independent experiments performed in triplicates with standard deviation (S.D). Unless indicated in figure legend, statistical analysis was performed using two-way ANOVA or unpaired t test (GraphPad Software, Inc., La Jolla, CA). The IC50 value was determined by nonlinear regression (curve fit) using log (inhibitor) versus response (three parameters) model.

---

**Supplemental Figure 1.**
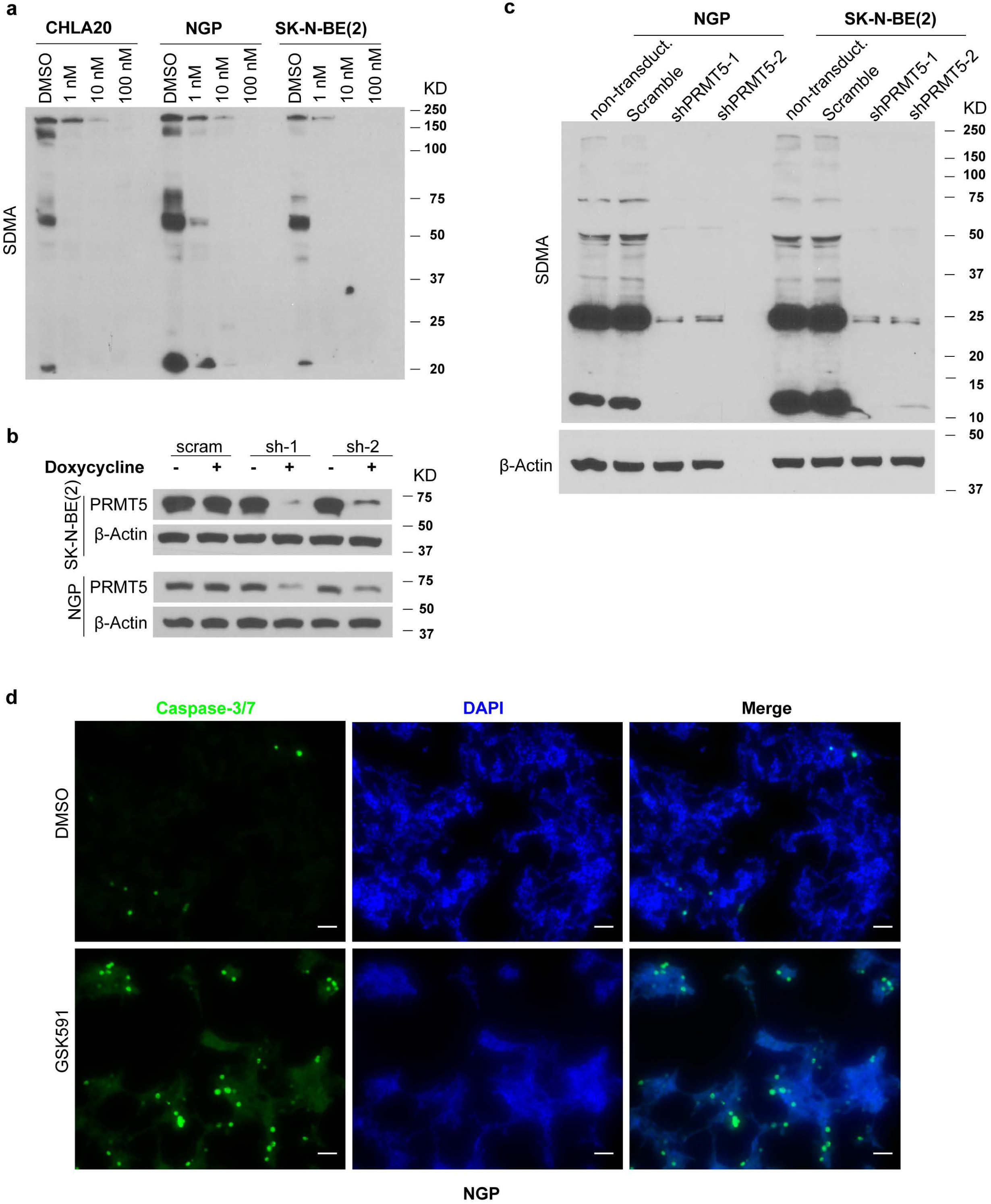
**a**, Immunoblotting of SDMA in neuroblastoma cells treated with DMSO or increasing doses of GSK 591. **b**, PRMT5 knockdown efficiency measured by Western blot in scramble or PRMT5 knockdown cells in the absence or presence of doxycycline. **c**, SDMA in control and PRMT5 knockdown cells. **d,** Caspase-3/7 staining in NGP cells treated with DMSO or GSK591. Scale bars, 100 *μ*m.

**Supplemental Figure 2.**
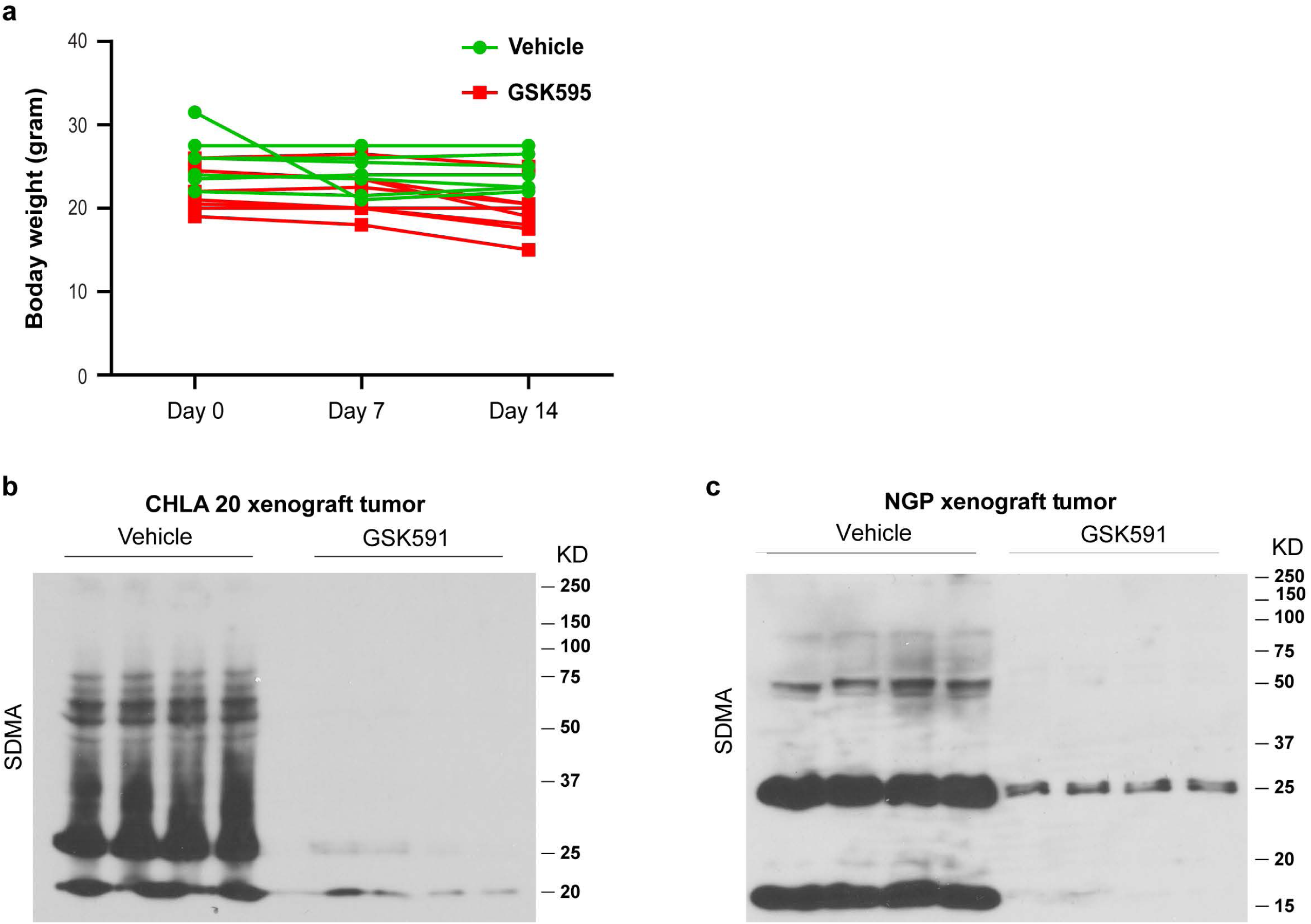
**a**, Weekly bodyweight of mice used in this study. Immunoblots of SDMA in CHLA20 (**b**) and NGP (**c**) xenograft tumors (equal loading control seen in Fig. 5d).

**Supplemental Figure 3.**
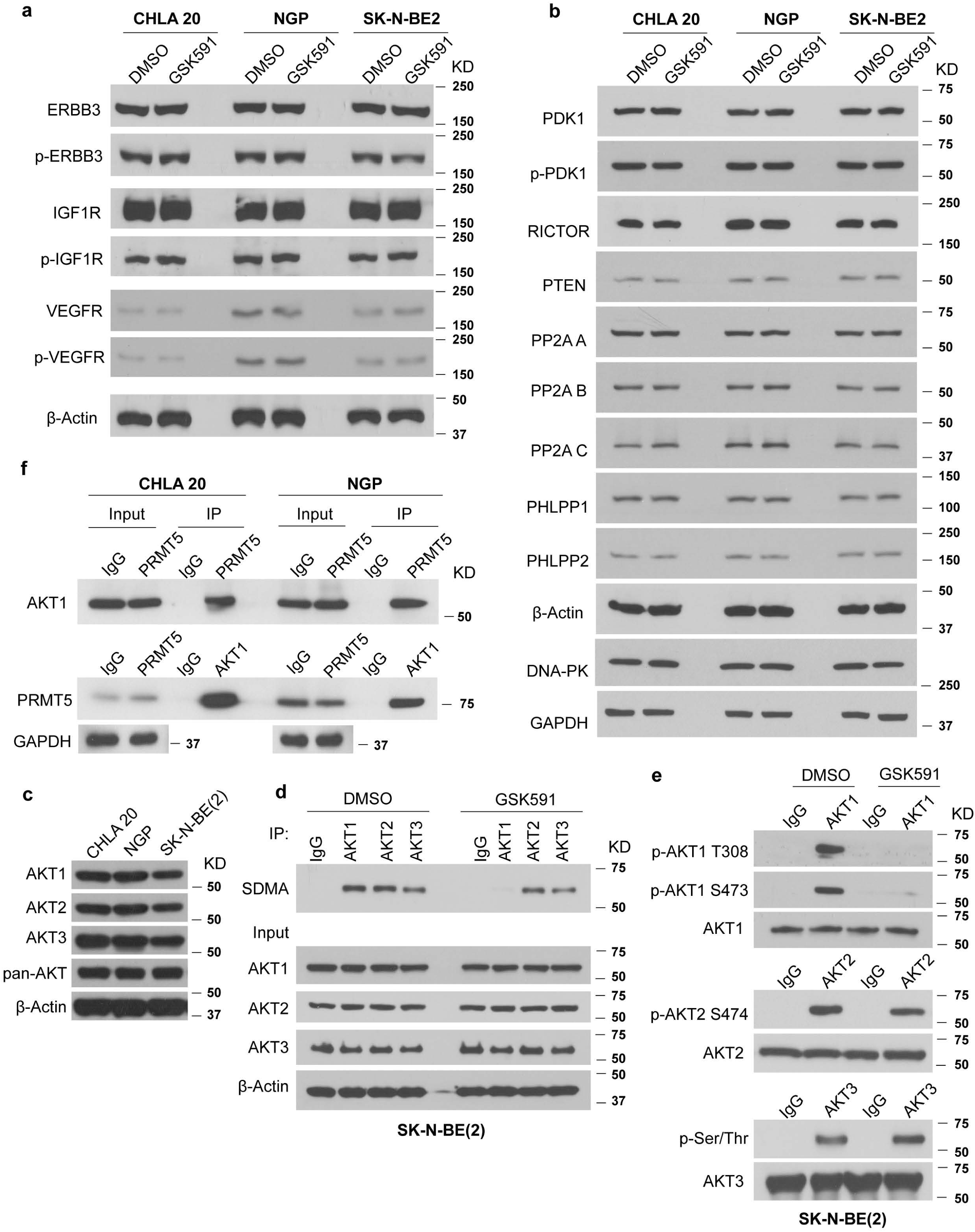
**a**, Immunoblots showing the protein levels or phosphorylation of RTKs not affected by GSK591 treatment. **b**, The expression or phosphorylation of kinases or phosphatases involved in the regulation of AKT activation in neuroblastoma cells under GSK591 treatment. **c**, The expression of AKT isoforms in neuroblastoma cell lines. **d**, SDMA of AKT isoforms in SK-N-BE (2) cells treated with DMSO or GSK591. **e**, Phosphorylation of AKT isoforms in SK-N-BE (2) cells upon GSK591 treatment. **f**, Western blots showing the interaction of AKT1 and PRMT5 by co-IP.

**Supplemental Figure 4.**
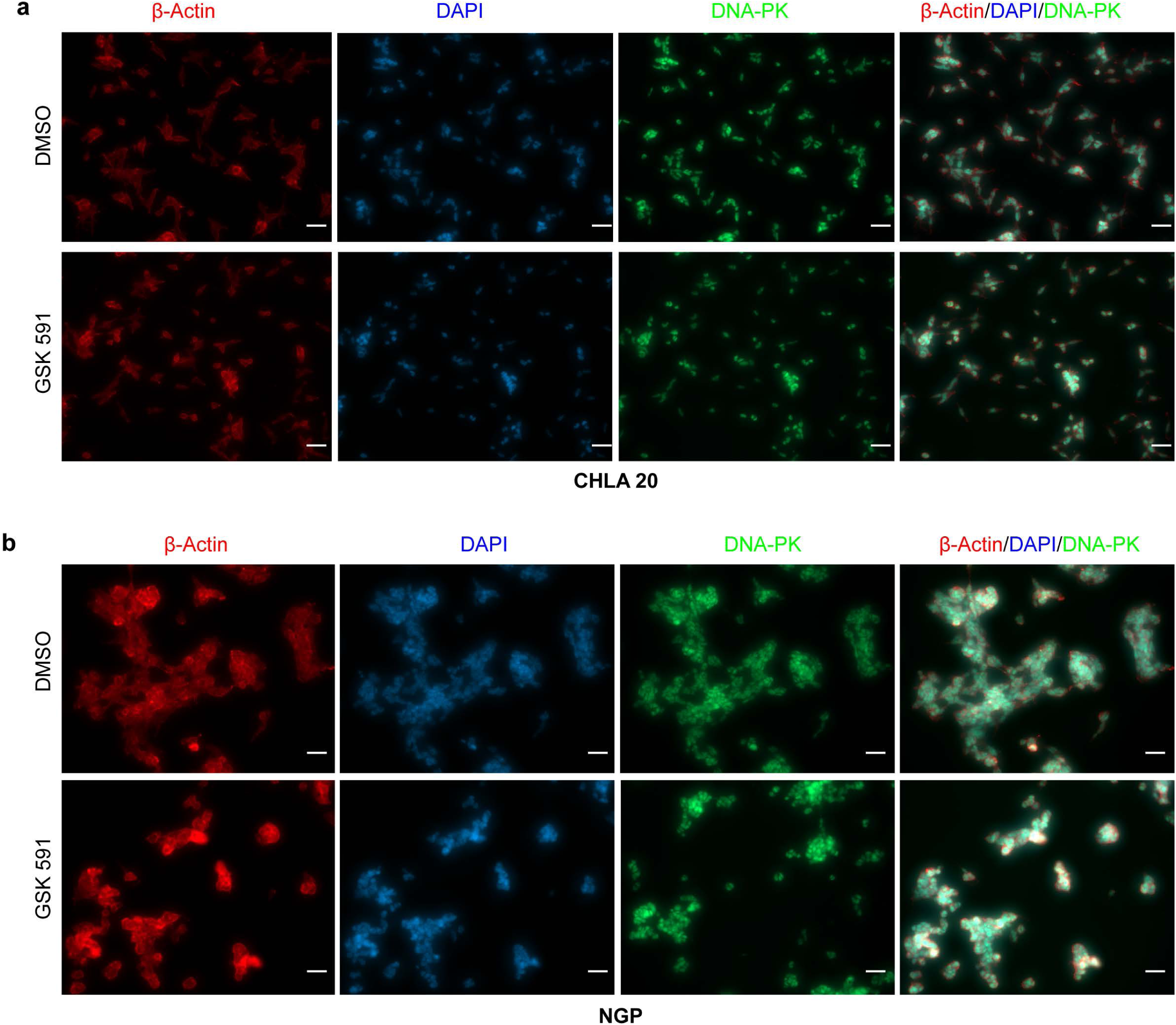
Immunofluorescence showing the expression of DNA-PKs in CHLA20 (**a**) and NGP (**b**) cells treated with DMSO or GS591. Scale bars, 100 *μ*m.

**Supplemental Figure 5.**
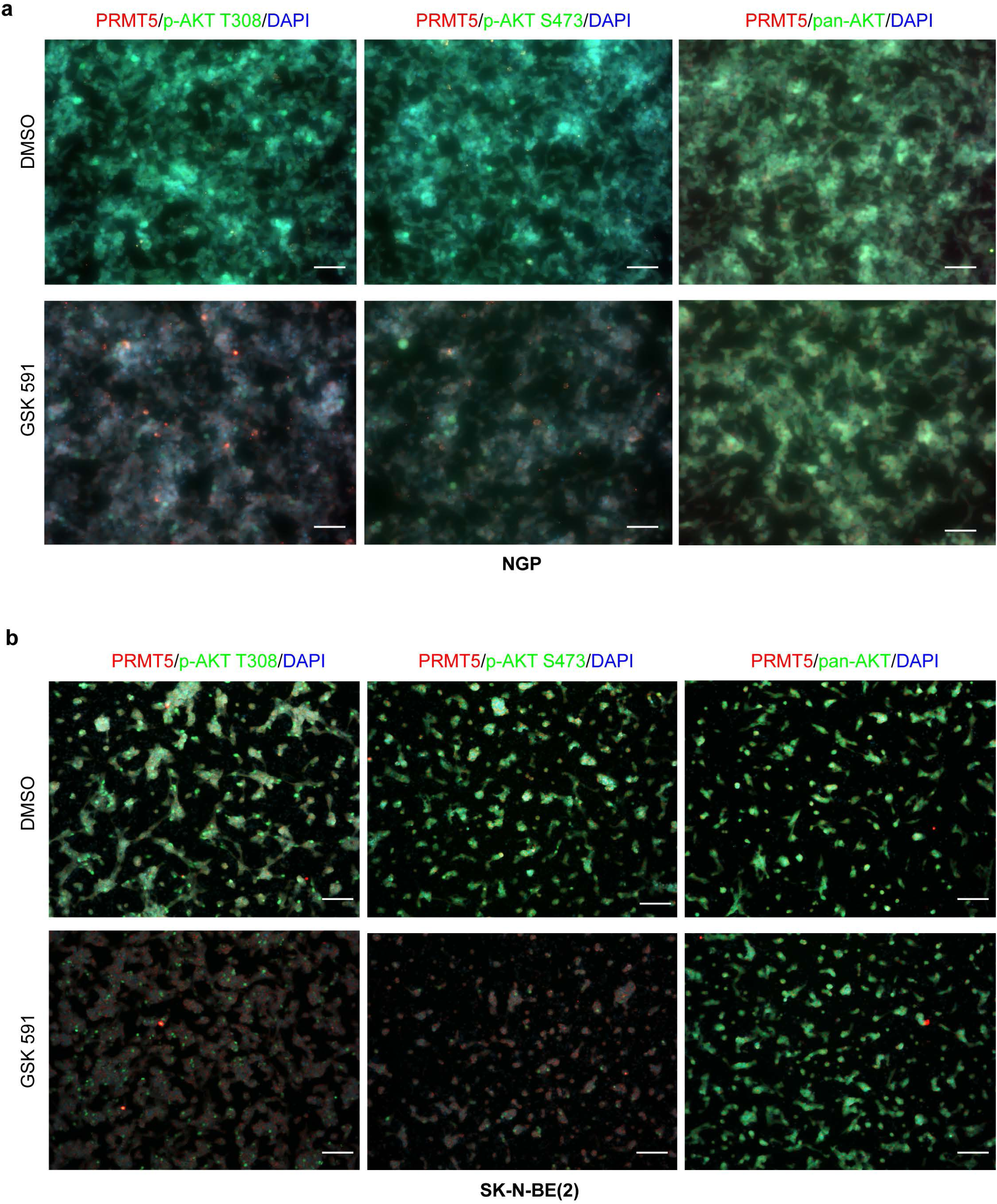
Co-localization of PRMT5 with p-AKT Thr308, p-AKT Ser473, and total AKT respectively by immunofluorescence in NGP (**a**) and SK-N-BE(2) (**b**) cells treated with DMSO or GSK591. Scale bars, 100 *μ*m.

